# Transcranial focused ultrasound induces source localizable cortical activation in resting state humans when applied concurrently with transcranial electric stimulation

**DOI:** 10.64898/2025.12.03.692212

**Authors:** Joshua Kosnoff, Colton Gonsisko, Kai Yu, Joseph Zhang, Yidan Ding, Yisha Zhang, Bin He

**Author notes:** Correspondence: Bin He, PhD.

## Abstract

The effects of transcranial focused ultrasound (tFUS) on the human brain are poorly understood. Currently, the field is at odds with whether tFUS is subthreshold modulating the brain’s excitability towards other stimuli, producing suprathreshold neural stimulation on its own, or if it even has a spatially specific non-auditory induced affect. Herein, we investigated the ability of tFUS, transcranial direct current stimulation (tDCS) and a novel combination of the two (transcranial electro-acoustic stimulation; tEAS) to evoke cortical target-location specific activity in 27 resting state humans with whole brain electroencephalography recordings. In none of the exogenous event related potentials did tFUS or tDCS result in location-specific activations. However, the co-modulatory combination of tEAS did, providing evidence that tFUS has a location-specific subthreshold modulatory effect. We propose a minimally modified Hodgkin Huxley model that explains our results and provides a unifying framework for the field-wide observed effects (or lack thereof) of tFUS.

## 1. INTRODUCTION

Therapeutic intervention of the human nervous system dates back centuries. Medical texts from c. 1^st^ century CE Rome^1^ explicitly prescribe uses of electric fish for pain management, and surviving records from c. 1550 BCE Egypt^2^ imply the practice may date even earlier. A modern, philosophical descendant of these methods, transcranial direct current stimulation (tDCS), applies subthreshold (meaning it will not induce neural firing on its own) current across brain tissue using non-invasive scalp electrodes^3^. The subthreshold current depolarizes (or hyperpolarizes, in the case of inhibitory current) the neurons towards (or away from) their action potential threshold, which will make the neurons more (or less) sensitive to additional stimuli. One of the biggest limitations with this form of neuromodulation has to do with the spatial precision at which it can be applied due to volume conduction effect^4^. Non-invasive electrical stimulation has spatial precision on the order of centimeters^5^, which can contain tens of millions of neurons^6^, and unintentionally modulate off-target brain regions as well^3,7^.

A recently introduced technique, transcranial focused ultrasound (tFUS), promises to overcome that limitation with a spatial precision on the order of millimeters^8,9^. tFUS’ mechanism of action continues to be under investigation, but one prevalent theory is the ultrasound pressure waves open mechanosensitive ion channels^10,11^. This theory would be consistent with behavioral outcomes dependent on the specific brain region targeted with tFUS, which have been reported in a range of human studies^8,12–17^.

However, there is a critical lack of understanding when it comes to tFUS’ effect on the human brain. The bulk of human tFUS studies pair it with a secondary stimulation (such as tactile vibrations^8,12^, transcranial magnetic stimulation (TMS) pulses^18^, or visual stimuli^15^), which point toward, but do not confirm, a subthreshold modulation-based mechanism of action. Other studies directly looked into tFUS’ ability to elicit percepts as the sole stimulus, but their results pointed towards, at best, an inconsistent ability to do so—a study investigating S1 activation only elicited percepts in 54% of trials^13^, and a study investigating V1 stimulation reported visual percepts in only 11 of 19 participants^14^. Beyond even the question of suprathreshold stimulation vs. subthreshold modulation, some studies report that the effects of tFUS may not actually be due to the pressure wave, but rather auditory conduction. tFUS, typically, is not applied as one continuous wave, but instead pulsed at some lower frequency (pulse repetition frequency; PRF; Figure 1d). This PRF makes a beeping noise, which is associated with activating auditory pathways^19,20^, and some studies go so far as to propose 100% of tFUS modulation is due to auditory activation^21,22^. In humans, it was reported that actively playing a tone over the duration of tFUS sonication led to no significant differences in EEG between sham and active conditions^23^.

**Figure 1:**
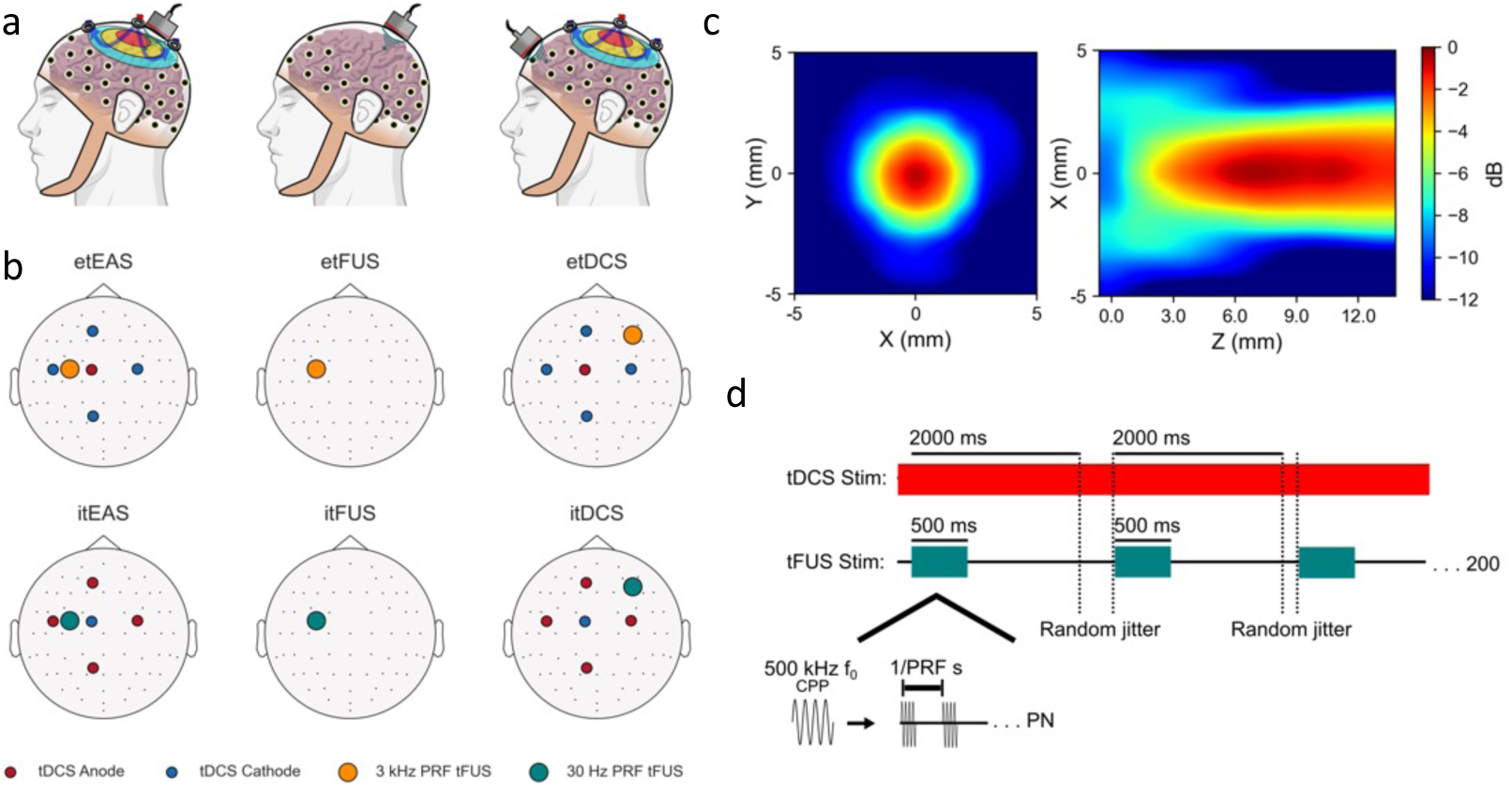
Methods. **a)** The experiment consisted of applying neuromodulation to eyes-closed resting-state humans. Transcranial electro-acoustic stimulation (tEAS) involved overlapping transcranial focused ultrasound (tFUS) and transcranial direct current stimulation (tDCS). The tFUS condition consisted of only tFUS. The tDCS control condition also involved a spatial control ultrasound, which did not overlap with the region enclosed by tDCS electrodes, to account for possible auditory confounds. **b)** A diagram of the experimental set up for each condition is provided with electrodes and transducers placed on their relative location to EEG electrodes. **c)** A free water scan of ultrasound is provided to illustrate its 5 mm full-width half-max (-6 dB) spatial specificity. **d)** For the experiment, 500 fundamental frequency (f0) ultrasound was pulsed every two seconds with up to an additional 20% random jitter for 200 trials. The pulse repetition frequencies (PRF) were either 30 Hz or 3 kHz, as specified. The cycles per pulse (CPP) and pulse number (PN) varied based on PRF to normalize all conditions to 500 ms sonication duration and a 30% duty cycle. For conditions involving tDCS, the electrical stimulation remained on throughout the entirety of the trial.

These studies raise critical questions about whether tFUS merely activates auditory pathways or if it evokes location-specific neuronal responses—and, if it does have a location-specific effect, whether that effect is subthreshold or suprathreshold. Resolving these questions requires comprehensive analysis of whole brain responses. While functional magnetic resonance imaging (fMRI) studies have investigated these questions and reported activations at the focal site of tFUS sonication^24,25^, their results may be confounded by multiple mechanisms. fMRI detects changes in blood-oxygen-level-dependent (BOLD) signal as a correlate for neural firing, but blood flow can be changed through a variety of methods not necessarily dictated by neuronal firing, such as vasodilation, which is directly inducible by ultrasound^26^. Further, unless tFUS is applied strictly parallel to the MRI’s magnetic field, physical interaction between the ultrasound pressure waves and the MRI’s magnetic fields will generate localized electric fields^27^, potentially creating artifactual neuronal activations absent in non-MRI environments. Consequently, fMRI measurements of tFUS’ neural activation ability are not definitive. Electroencephalogram (EEG) based recordings, which directly measure electrical activity without the need for magnetic fields, may offer more insight into the true effects of tFUS as a sole stimulus.

Herein, we apply excitatory (e) and inhibitory (i) tDCS, tFUS (500 kHz fundamental frequency), and a novel combination of the modalities (transcranial electro-acoustic stimulation; tEAS) to the cortex of resting-state humans (Figure 1). In follow-up studies, we broaden our tFUS parameter scope. We record and analyze simultaneous whole-scalp EEG recordings and image their corresponding cortical sources to characterize targeted-location specific, or lack-thereof, responses. We also present a first-principles verification of our findings using a minimally modified Hodgkin Huxley model.

## RESULTS

### Transcranial electro-acoustic stimulation produces a hemisphere specific exogenous response

Butterfly plots of each condition’s preprocessed EEG are shown in Figure 2a. Statistical differences between EEG signals of the region of interest (ROI; electrodes C1, C3, FC1, and FC3) and their contralateral homologous counterparts (C2, C4, FC2, and FC4) were assessed using spatio-temporal permutation cluster tests within the 0 – 200 ms exogenous (stimulus-evoked) response window. The tests identified significant differences (*p* < 0.05) in the itEAS (30 Hz PRF tFUS with overlapping cathodal tDCS) and etEAS (3 kHz PRF tFUS with overlapping anodal tDCS) conditions, but none for itFUS (30 Hz PRF), etFUS (3 kHz PRF), itDCS (cathodal tDCS with spatial-control 30 Hz PRF tFUS) or etDCS (anodal tDCS with spatial-control 3 kHz PRF tFUS) conditions (Figure 2b). Both significant clusters encompassed electrode C3 and the 10 – 63 ms time window. We used the shared components across both tEAS conditions’ significant clusters as spatial and temporal windows for further analysis.

**Figure 2:**
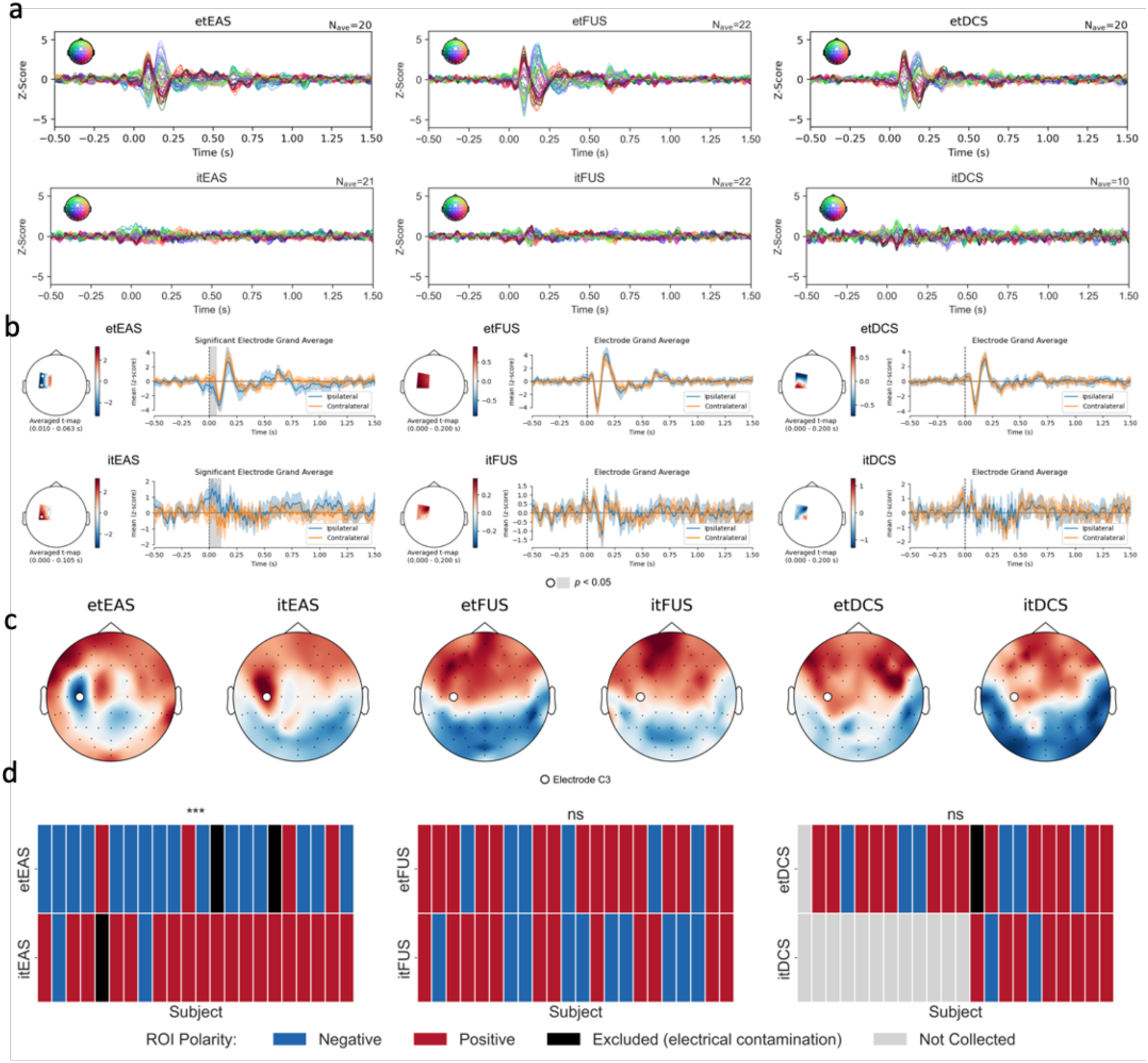
Transcranial electro-acoustic stimulation (tEAS) induces an exogenous hemisphere-specific response. **a)** Preprocessed EEG butterfly plots of the experiment show exogenous (< 200 ms post stimulation) event-related potentials at 30, 90 and 170 ms for conditions with 3 kHz PRF tFUS**. b)** Spatio-temporal paired t-test permutation cluster tests were performed comparing the ipsilateral ROI electrodes (C1, C3, FC1, and FC3) against their homologous contralateral counterparts (C2, C4, FC2, and FC4) for the 0 to 200 ms window. Significant electrodes are marked in with white circles on the right subplot t-statistic topographic maps and significant corresponding time windows are shaded in gray on the left subplot. The temporal traces (mean ± 95% confidence interval) are a grand average of the significant electrodes, or, if there are no significant clusters, of all the electrodes. Only the tEAS conditions yielded significant clusters, indicating the 3 kHz PRF ERPs are symmetrically distributed across the brain. The largest overlapping time window between the significant clusters, 10 – 63 ms, will be used for further analysis. **c)** The 10 – 63 ms topographic maps visually indicate a strong ROI response in i/etEAS conditions. Electrode C3 is marked with a white circle. **d)** Electrode C3 was identified in both significant clusters from tEAS permutation cluster tests. Further statistical analysis (two-tailed McNamar’s paired binary data test with fdr multiple comparison correction) was conducted to test for a significant effect on C3 polarity between e/i conditions for all modulation parameters. i/etEAS was found to have a significant effect (*padj* < 0.001). No significant effects on polarity were detected for tFUS or tDCS.

### Inhibitory vs. excitatory tEAS produces significant changes to EEG polarity

EEG data were averaged over the permutation-cluster test identified time frame of 10 – 63 ms. Resultant topographic maps present with visually apparent ROI activations for tEAS conditions and primarily hemisphere-symmetric topographies for the other conditions (Figure 2c). Effects on electrode C3 (selected based on its presence in permutation cluster-test results) polarity for this time average were assessed with McNemar’s test and False Discovery Rate (FDR) multiple comparison correction (Figure 2d). Inhibitory tEAS led to a significant change (*p_adj_* < 0.001) in ROI polarity compared to etEAS. Neither e/i tFUS nor the e/i tDCS conditions induced significant polarity changes (*p_adj_* > 0.05).

### tEAS induces significantly localizable responses to the targeted ROI

EEG data were back projected to the subjects’ cortices by source localization using the dynamic statistical parametric mapping (dSPM) method, and the resultant power of source waveforms were averaged over the 10 – 63 ms window. Source-morphing to a common brain model and averaging the response across subjects shows strong concordance with the targeted ROI, defined by the subspace contained within electrodes C1, C3, FC1, and FC3 orthogonally projected to the cortex (Figure 3a). Statistical analysis to determine if ipsilateral (IL) ROI power was significantly higher than the homologous contralateral (CL) power was conducted with permutation t-tests with FDR multiple comparison correction. Both tEAS conditions had significantly increased IL ROI source power (etEAS IL = 2.61 ± 1.21 z_dSPM_, CL = 1.66 ± 0.66 z_dSPM_, *p_adj_* < 0.01; itEAS IL =2.39 ± 1.14 z_dSPM_, CL = 1.20 ± 0.47 z_dSPM_, *p_adj_* < 0.01). None of the other conditions had significantly increased IL source power (*p_adj_* > 0.05; Figure 3b).

**Figure 3:**
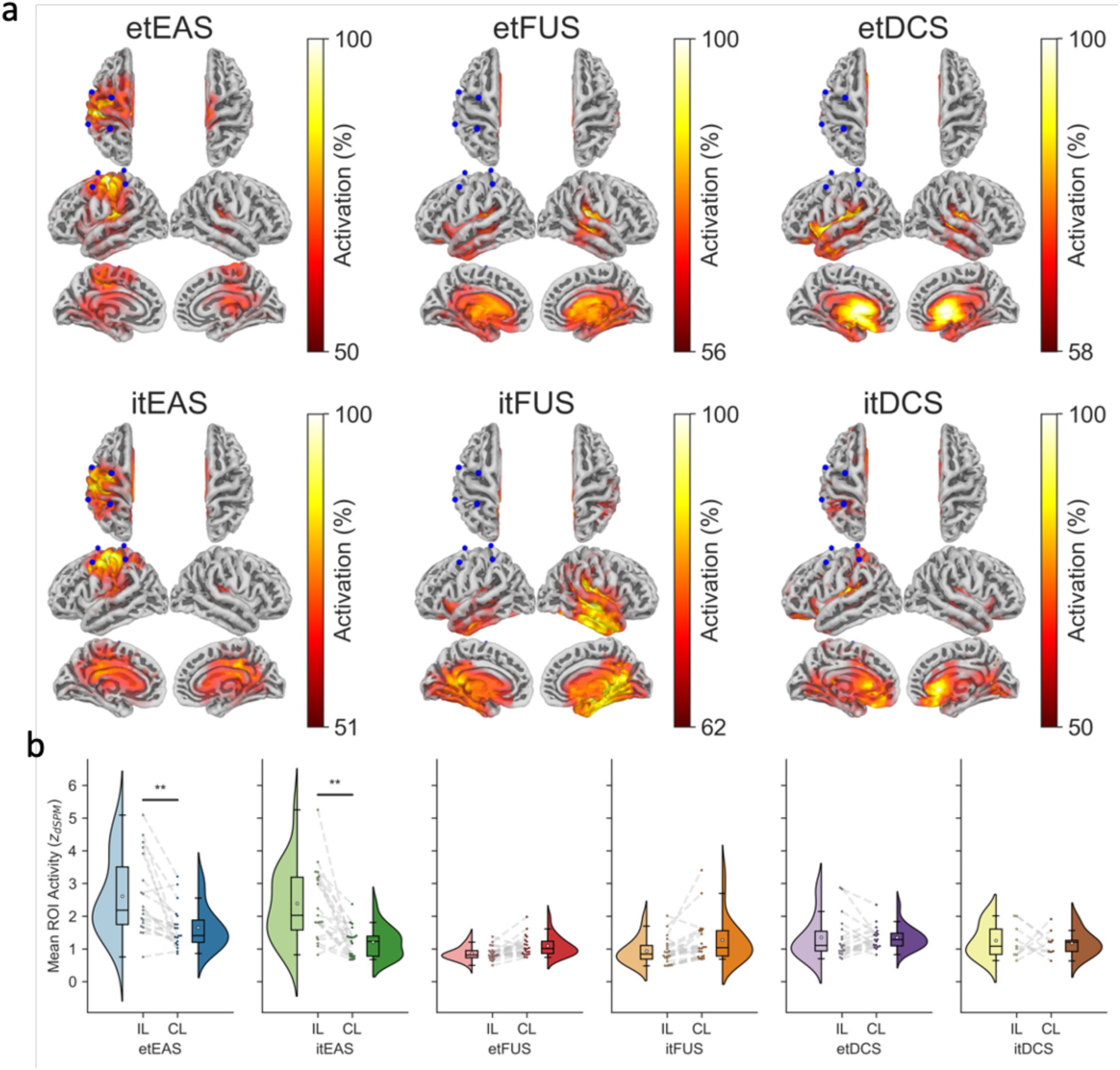
tEAS induces source localizable cortical responses. **a)** EEG data were projected to the cortex using the dSPM source imaging method. Subject data were averaged over the 10 – 63 ms post-stimulation window and source morphed to FreeSurfer’s *FSAverage* common brain model and averaged together for visualization. The cut-off level of activity for brain sources were defined based on Otsu’s threshold. **b)** i/etEAS led to significantly higher ipsilateral ROI source activity than contralateral ROI source activity (one-tailed t-test with fdr multiple comparison correction; \*\**padj* < 0.01). e/itFUS and e/itDCS did not (*padj* > 0.05).

Pairwise comparisons using each subject’s condition average were made between IL source power, as well as precision, recall, and F1 score of source localization with respect to the ROI (Supplemental Figure S1). For all metrics, tEAS conditions were found to have significantly higher outcomes. Cohen’s *d* analysis yielded that tEAS conditions had at least moderate effect sizes (*d* > 0.5) for all comparisons with other conditions, and often times large effects (*d* > 0.8). Pairwise FDR-adjusted *p-*values and Cohen’s *d* values are provided within the relevant sub-figures.

### Transcranial focused ultrasound results in significant auditory activations

Despite the presence of three distinct exogenous time window ERPs induced from 3 kHz tFUS in the topographic maps (30, 90, and 170 ms peaks; Figure 2a), spatio-temporal permutation cluster tests did not return any significant differences for left hemisphere ROI electrodes compared to their right hemisphere counterparts. This points to a hemisphere-symmetric response, despite tFUS only being applied to one side. Topographic map visualizations of each peak (20 – 40 ms, 80 – 100 ms, and 160 – 180 ms averages; Figure 4a) do not provide any evidence of a target-location specific effect. Source imaging metrics of these peak windows continue to support a lack of ROI response (20 – 40 ms precision w.r.t. ROI: 0.013 ± 0.014; 80 – 100 ms: 0.015 ± 0.019; 160 – 180 ms: 0.022 ± 0.020).

**Figure 4:**
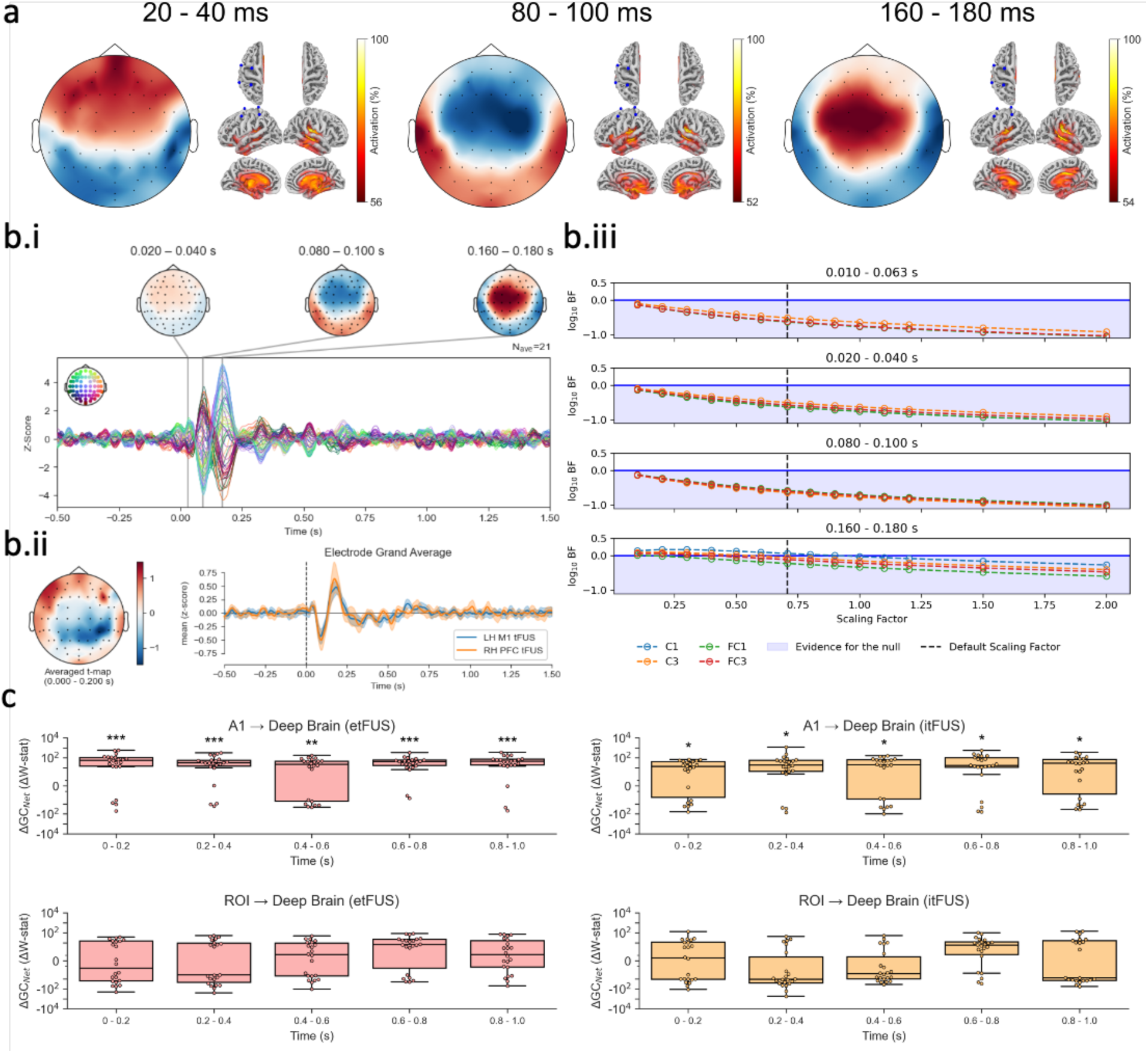
tFUS induces significant auditory activations. In all three conditions with 3 kHz tFUS (etFUS, etDCS, and etEAS), distinct ERPs at 30, 90, and 170 ms were apparent (Figure 2a). **a)** For etFUS, EEG and dSPM source activities were averaged for ±10 ms around these ERPs to elucidate their origin. At all three time windows, the topographic maps present with symmetric responses across hemispheres. Despite tFUS being targeted to the left primary motor cortex area, source imaging of the peaks show the underlying sources are located in the auditory or deep brain. **b)** The electrophysiological response of tFUS applied to resting-state left motor cortex is indistinguishable from tFUS applied to resting-state prefrontal cortex **b.i)** In 21 of the subjects, a 3kHz spatial control was conducted where tFUS was applied to the right prefrontal cortex (PFC). Even though tFUS was applied contralaterally and to another region, the resultant waveform and ERP spatial distributions are strikingly similar to tFUS applied at M1. **b.ii)** Spatiotemporal cluster testing did not return any significant clusters between tFUS to left hemisphere (LH) M1 and tFUS to right hemisphere (RH) PRF. **b.iii)** The lack of significant differences through traditional hypothesis testing methods does not indicate statistical similarity. However, using Bayes Factor (BF) analysis, the evidence in favor of the null (i.e., that two distributions are statistically the same) can be quantified. A log10 BF < 0 is evidence for the null, whereas log10 BF > 0.5 is considered evidence against the null. To address BF analysis’ sensitivity to its scaling factor (the commonly selected default value is 0.707), we ran BF analysis for RH PFC tFUS vs. LH M1 tFUS ROI electrodes over a wide range of values (0.1 – 2.0). In the 10 – 63, 20 – 40, and 80 – 100 ms windows, for every scaling factor tested, all four electrodes resulted in negative log10 BFs, providing robust evidence that the ROI response is the same across these two conditions. For C1, C3, and FC3 in the 160 – 180 ms window, BF analysis was inconclusive for possibly small-medium effect sizes (scaling factor < 0.707; 0 < log10 BF < 0.5), though it continued to provide robust evidence in favor of null for medium-large effect sizes (scaling factor > 0.707). **c)** Granger Causality analysis reveals significant tFUS-induced auditory cortex to deep brain projections. The Net Granger Causality (GCNet) Wald statistic was calculated for each time bin by subtracting the GC of the reverse flow from the GC of the forward flow of information of auditory cortex to the deep brain and the left M1 ROI to the left deep brain for both etFUS and itFUS. The GCNet values were normalized by subtracting the GCNet for the baseline period (-0.5 to -0.1 s) for each subject’s pathways (ΔGCNet). For auditory to deep brain activations, the cortical hemisphere with the maximum net flow was considered in analysis for each subject. For M1, only the tFUS-targeted left hemisphere ROI to left deep brain parcellation was considered. The differences between the baseline GCNet and each time bin’s GCNet were analyzed with permutation t-tests. FDR multiple comparison correction was applied locally. Both etFUS and itFUS induced significant auditory to deep brain Granger causality, while neither induced a significant ROI to deep brain flow of information. \**padj* < 0.05, \*\**padj* < 0.01, \*\*\**padj* < 0.001.

In 21 of the 22 subjects with etFUS, we also tested a 3 kHz PRF right prefrontal cortex (RH PFC) spatial control condition. The resultant waveforms and ERP spatial topographies (Figure 4b.i) share a striking resemblance to tFUS applied to left motor cortex area (LH M1). Spatio-temporal permutation cluster testing did not yield any significantly different clusters between LH M1 and RH PFC conditions (Figure 4b.ii). Further, when we directly quantified statistical similarity between the LH ROI electrodes for the two conditions using Bayes Factor (BF) analysis, the evidence that the 10 – 63, 20 – 40, and 80 – 100 ms responses of the two conditions were the same was robust (log_10_ BF < 0 for all electrodes/scaling factors; Figure 4b.iii). For the 160 – 180 ms window, BF analysis was inconclusive for small-medium effect sizes (scaling factors < 0.707; 0 < log_10_ BF < 0.5), though it continued to provide evidence in favor of similarity for medium-large effect sizes (scaling factors > 0.707). For comparison, BF analysis of etEAS compared to etFUS yielded evidence that the certain ROI electrode responses were significantly different (log_10_ BF > 0.5; Supplemental Figure S2).

Granger causality analysis of both itFUS and etFUS conditions shows significant increases in information flow from primary auditory cortex to the deep brain, and no changes in net information flow from the targeted ROI to the deep brain (Figure 4c). In a follow-up study with 11 subjects, it was found that even an inaudible PRF (30 kHz) continued to induce significant auditory to deep brain Granger causality (Supplemental Figure S3).

tEAS conditions were also found to induce significant increases in auditory to deep brain connectivity, and have no effect on the net flow of information from the targeted ROI to deep brain (Supplemental Figure S4).

### tFUS, but not tEAS, induces endogenous dissociations between the ROI and deep brain

Despite not inducing a significant effect on net Granger causality, tFUS was found to have significant dissociation effects on undirected spectral imaginary coherence (ImCoh; Supplemental Figure S5). Both itFUS and etFUS significantly reduced the gamma band ImCoh in endogenous (> 200 ms) time windows. etFUS also significantly dissociated alpha band connectivity. None of these dissociations were present in the tEAS conditions.

### Increased tFUS parameter searches continued to yield no evidence of ROI suprathreshold stimulation

A follow-up call-back study was conducted in 10 of the 22 original subjects, in which additional tFUS parameters were tested. These parameters included 1,500 and 30 Hz PRFs with 30% duty cycle (DC), and, initially, 10% and 60% DC with both 30 and 3,000 Hz PRFs. All four of the first four subjects experienced thermal discomfort with 60% DC, and so we did not test it further. From the fourth subject onward, we tested 0.6% DC instead. Source imaging results continued to point towards deep brain and temporal lobe activations (Figure 5a) and ROI analysis continued to find no increases in ipsilateral source power for any of the new tested conditions (Figure 5b). Analysis of the effects of holding the DC or PRF constant and changing the other parameter yielded no significant differences (permutation one-way ANOVA; all *p* > 0.05) and Bayes Factors provide robust evidence that the responses across conditions are statistically similar (log_10_ BF < 0; Figure 5c).

**Figure 5:**
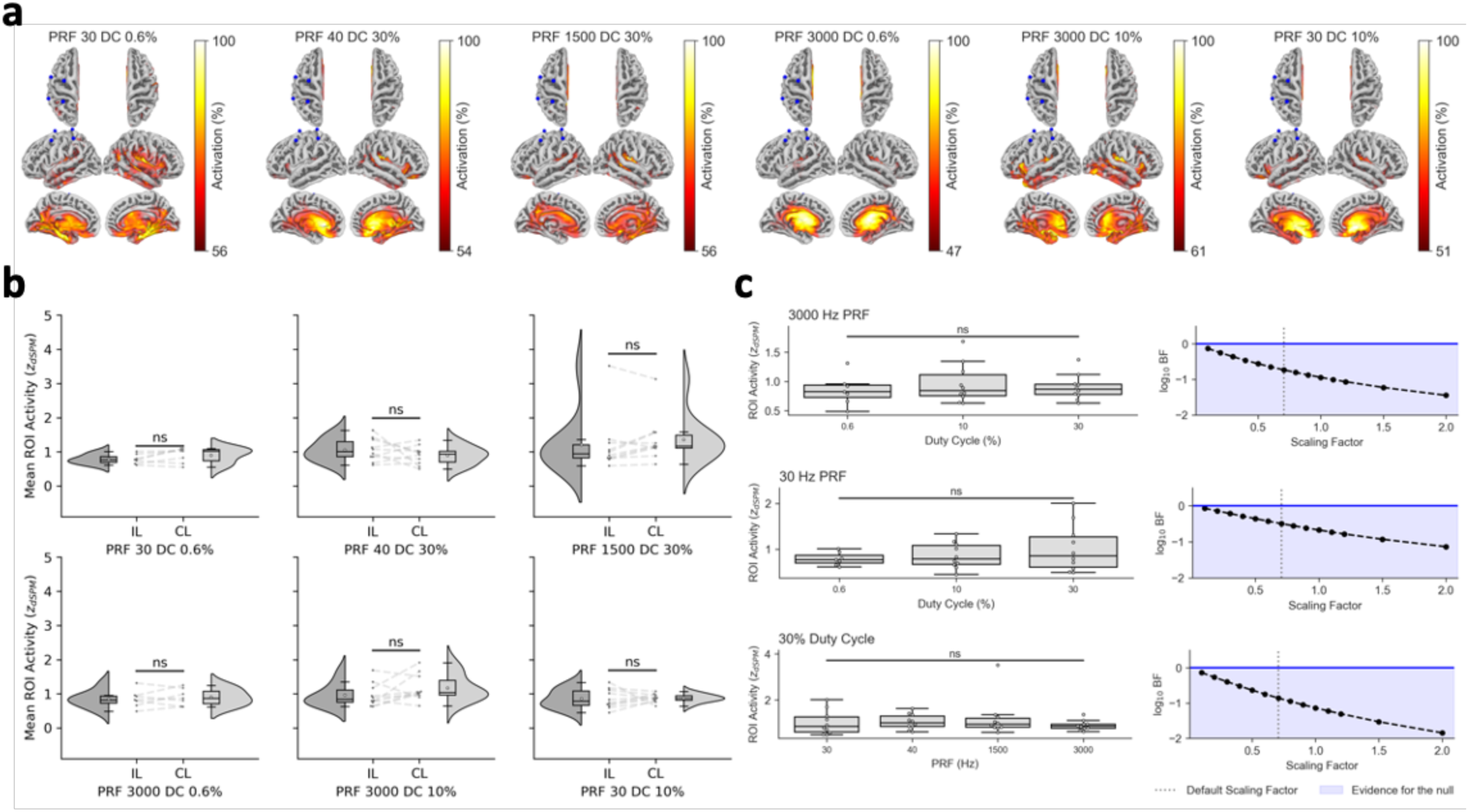
An expanded parameter search for tFUS continues to find no source localizable activity in resting-state humans. A follow-up study was conducted in 10 subjects to test for possible changes in activity from varying the duty cycle or PRF from the original study parameters. **a)** EEG data were projected to the cortex using the dSPM source imaging method. Subject data were averaged over the 10 – 63 ms post-stimulation window and source morphed to FreeSurfer’s *FSAverage* common brain model and averaged together for visualization. The cut-off level of activity for brain sources were defined based on Otsu’s threshold. Visualizations continue to show strong deep brain and temporal lobe activations, but nothing concordant with the ROI. **b**) No tFUS-only parameter combination yielded significantly higher ipsilateral than ROI source activity (one-tailed permutation t-test with FDR multiple comparison correction; all *padj* > 0.05). **c)** Comparisons across the subjects’ tFUS-only ipsilateral ROI responses were compared for constant PRFs and a constant Duty Cycle. Their 30 Hz and 3 kHz PRF with 30% DC data from the initial session were also included in this analysis. No significant changes were detected with a permutation one-way ANOVA test (all *padj* > 0.05). Beyond the lack of statistic differences, Bayes Factor analysis indicates the resting-state tFUS responses are statistically similar across the parameter spaces.

In an additional follow-up study with 11 subjects, the effects of higher pressure were investigated by normalizing the *in situ* pressure to 300 kPa with Sim4Life simulations (original study *in situ* mean pressure was 231 kPa). The sonication duration was reduced from 500 ms to 100 ms to compensate for this increase. Even still, 2 of the 11 subjects (18.2%) experienced scalp thermal discomfort and their experiments were aborted. Analysis of the other 9 subjects’ data continued to yield no significant increases in IL compared to CL ROI (Supplemental Figure S6; *p* > 0.05).

### tEAS is more than a linear sum of its parts

We took the summation of the source-space tFUS and tDCS conditions to produce a condition representing the sum of independent electric and acoustic stimulation. The summation did not result in increased concordance with the ROI (Supplemental Figure S7a), nor to significant increases in ipsilateral over contralateral ROI source power (Supplemental Figure S7b; permutation t-test *p* > 0.05).

### tEAS response is driven by tDCS polarity, not tFUS PRF

Anodal tDCS was combined with 30 and 3,000 Hz PRF tFUS and analyzed over the 10 – 63 ms window (Supplemental Figure S8). Both conditions led to brain sources concordant with the ROI. Comparisons of the ROI source power yielded no significant differences across conditions (permutation t-test *p* > 0.05), and Bayes Factor analysis provided evidence that the response powers were statistically similar (log_10_ BF < 0).

### A minimally modified Hodgkin-Huxley model provides support for an electro-acoustic co-modulatory effect

The Hodgkin-Huxley model was modified to have an additional parallel pathway representing piezo-electric ion channels (Figure 6a). Simulations revealed that subthreshold ultrasound-based opening of piezo-electric channels (Figure 6b) and subthreshold direct current, representing tDCS (Figure 6c), could combine to form a super-threshold stimulus (Figure 6d-f).

**Figure 6:**
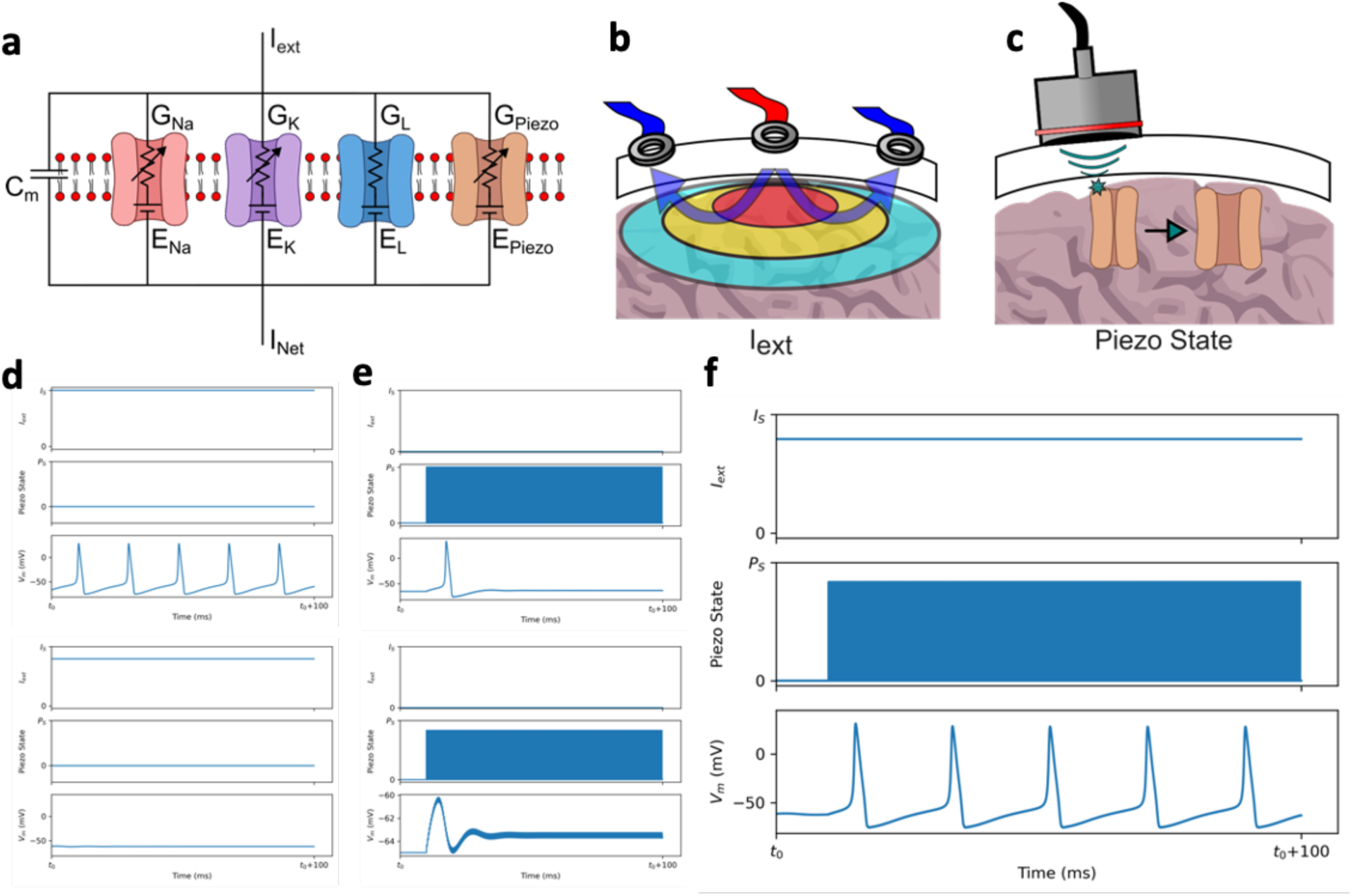
A minimally-modified Hodgkin-Huxley model can help explain the electro-acoustic effect. **a)** The Hodgkin-Huxley model was expanded to include a piezo-electric channel. **b)** tDCS drives electricity across the brain, which is an externally applied current. **c)** tFUS can mechanically open or close piezo-electric channels, the effect of which can be modeled as a change in the probability that the piezo-electric channel is open (Piezo State). **d)** (Top) There exists some level of current, IS, which will cause the neuron to produce action potentials. (Bottom) Below that threshold, applied current will not produce action potentials. **e)** (Top) there exists some Piezo State, PS, that will lead to an action potential. (Bottom) A probability below that threshold will not lead to an action potential. **f)** A sub-threshold applied external current and sub-threshold piezo-electric channel probability can combine to produce action potentials.

## 2. DISCUSSION

While the effect of tFUS is commonly considered to be based on the targeted brain region^8,9,12–17,28^, there is a lack of agreement for whether tFUS is modulating neural excitability or stimulating the neurons on its own. Beyond that, recent discoveries in tFUS research have suggested that auditory pathways, but not the focused ultrasound *per se*, are responsible for modulating brain signals^21–23,29^. We investigated this phenomenon by applying tFUS, tDCS, and a co-modulatory combination of the two, tEAS, to resting-state humans and characterizing the electrophysiological responses with whole brain EEG and EEG source imaging techniques. In follow-up studies, we broadened our tFUS parameter search to include additional PRFs, duty cycles, and higher *in situ* pressure. Our results indicated that, while neither resting-state tFUS nor tDCS produced targeted location EEG source localizable brain activities on their own, when they were applied together as tEAS, the multi-modal co-modulatory combination could. Our data provide strong evidence that, while there are prominent auditory effects, the applied tFUS has location-specific subthreshold modulatory effects on the human cortex.

When the ipsilateral ROI electrodes’ activities were compared against the contralateral homologous counterparts, only the co-modulatory tEAS conditions yielded significant differences (Figure 2b). While the clusters from permutation cluster tests are not guaranteed to be significant for their entire extent^30^, we did consider the overlapping windows (10 – 63 ms) for significant itEAS and etEAS clusters to be interesting enough due to their shared presence to warrant use in our further analysis. We continued our analysis in the cortical domain using dSPM EEG source imaging. The subject average response had strong concordance with the targeted ROI for both itEAS and etEAS, but not for the other conditions (Figure 3a). The ipsilateral ROI source power was significantly higher than that of the homologous contralateral ROI for only the tEAS conditions (Figure 3b). In a follow-up study, we expanded our resting-state tFUS parameter search to include additional 40 and 1,500 Hz PRFs with 30% DC, as well as 0.6%, 10%, and 60% DCs with both 30 and 3,000 Hz PRFs. 60% DC led to unpleasant scalp sensations^31^ in all initial (4/4) subjects and the experiments were aborted. Analysis of the other parameters continued to show a lack of sonication-location specific suprathreshold responses (Figure 5).

Directly comparing ipsilateral ROI source power and source localization classification metrics (Precision, Recall, and F1 Score) across conditions indicated tEAS had significantly higher ipsilateral outcomes compared to the others (Supplemental Figure S1). Summing the separate tDCS and tFUS conditions together did not yield spatially specific information (Supplemental Figure S7), providing additional evidence for a compounded electro-acoustic effect.

Not only did itEAS and etEAS induce significantly higher source localization metrics and source power than the other tested conditions, but they also led to a significant change in ROI polarity (Figure 2d). While this polarity change is evidence of differing electrophysiological effects, a follow-up study indicates that this change in response is driven from tDCS polarity and not tFUS PRF (Supplemental Figure S7). Anodal tEAS experiments were repeated with both 30 Hz and 3 kHz PRF tFUS and the resultant ROI responses were found to be statistically similar through Bayes Factor analysis. This is consistent with another research group’s report that neural response may not be a function of PRF^32^. Our prior human research has indicated tFUS may act on feature-based attention^16^, so it may also be possible that PRF-dependent responses would be brought out with non-resting-state task engagement. Understanding the translation from animal models to human models remains a critical challenge in neuroscience work, and additional research into neuron response to tFUS PRF at the human brain scale is warranted.

Despite a lack of source localizable activity and changes to granger causality net information flow, both etFUS and itFUS were found to have significantly reduced spectral connectivity from ipsilateral ROI to deep brain compared to the contralateral ROI to deep brain (Supplemental Figure S5). This is consistent with other literature that have reported tFUS to have a target-location specific dissociation effects^24,33^. etFUS and itFUS significantly modulated different time windows of gamma band connectivity, and only etFUS significantly modulated the lpha band, providing some evidence of circuit-level PRF-dependencies. However, there were no spectral connectivity changes present in the tEAS conditions. At first, this seems counter intuitive, as one would expect the suprathreshold tEAS condition to have stronger, compounded effects on brain connectivity changes. Instead, our data indicate that tEAS overrides the spectral dissociation tFUS otherwise induces. The timings of these connectivity changes point to why this might be the case. All but one of the spectral connectivity changes occur after 200 ms of sonication onset. The first 200 ms of brain activity post-stimulation is often considered exogenous, or stimulus driven, while activity occurring later than that is considered endogenous, and is reflective of internal cognitive processing^34^. Given this, these dissociations are likely a result of tFUS inducing some internal change in cognitive brain-state, which can be overridden with suprathreshold stimulation. Additional research is warranted on the nature and function of tFUS’ dissociations.

Excitatory (3 kHz PRF) tFUS targeted to the left motor region, as part of etFUS, etEAS, and etDCS, induced distinct ERPs at approximately 30, 90, and 170 ms post stimulation onset (Figure 2a, Figure 4a). This is consistent with a previous study that identified V1-targeted tFUS induced EEG peaks, though their reported peak times were labeled as 55 ms, 100 ms, and 150 ms post sonication^14^. However, when we compared these ERPs to those induced by applying tFUS to an entirely different brain region (right hemisphere prefrontal cortex), the resultant waveforms did not just lack statistical differences from each other—they were found to be statistically similar through Bayes Factor analysis (Figure 4b). In addition, Granger causality investigations revealed significantly increased A1 to deep brain information flow for both 30 Hz and 3 kHz tFUS conditions (Figure 4c). These results provide support on a possible auditory based effect of tFUS neuromodulation in humans^21–23,29^. Indeed, the 3 kHz PRF EEG topographic spatial distributions of the 90 and 170 ms ERPs do share visual resemblance to the previously reported auditory induced N1 and P2 ERPs^23^. Nevertheless, the present results that combining tFUS with tDCS led to spatially specific responses, when neither could do so individually (Figure 3), as well as tFUS dissociating the spectral connectivity between the ROI and the deep brain (Supplemental Figure S5), indicate that tFUS does have targeted-location specific effects.

A minimally modified Hodgkin-Huxley (HH) model sheds more insight into what might be occurring (Figure 6). We modeled the electro-acoustic effect with an additional piezo-electric ion channel within the traditional HH neuron. It supported that subthreshold current can be combined with subthreshold ultrasound pressure to induce a suprathreshold response. Since the model was created with parameters calculated on different species (the original HH parameters are from a squid^35^, piezo-electric components are from rodent measurements^36–38^, and the model is currently being used to test findings in humans), particular values of the current or piezo-electric channel open probability needed to produce spikes likely do not hold much meaning to any particular species. However, the model does provide a high-level unifying framework on the previously reported effects of tFUS.

In most of the published literature, tFUS is used explicitly in conjunction with another stimulus. These published paired stimuli include tactile vibration^8,12^, physical movement^17^, transcranial magnetic stimulation^18^, visual stimulation^15,16^, fMRI (which can introduces artifacts mentioned in the introduction)^24,25,27^, and now, as part of this study, tDCS. Even in the studies that investigated the effects of tFUS to induce tactile^13^ or visual^14^ perceptions, tFUS likely co-interacted with an attention-modulated somatosensory or vision circuity, as the participants were instructed to describe the location and type of perceived sensations, which could put their brains into attended states. This is further supported by another ultrasound-to-visual cortex perception study that found even the sham condition led to visual hallucinations^39^. Indeed, taking all of this into account within the context of our findings, it does imply that tFUS may be a subthreshold co-modulator, and not a suprathreshold stimulator. This may be why some researchers^21–23^, present study included, have not found tFUS to have a non-auditory related source localizable effect from EEG when not paired with another stimulus.

At the same time, the model supports that there is some open probability of piezo-electric channels, and thereby ultrasound pressure level, that will induce neuronal firing. This is in line with rodent cortex studies that have reported tFUS’ ability to be the sole stimulus^9^, and induce source localizable effects^40^. To further investigate this in humans, we repeated our experiment on a cohort of 11 subjects, increasing the target pressure to a normalized 300 kPa using Sim4Life simulations (Supplemental Figure S6). To lower the possibility of the transducer overheating, we reduced the sonication duration from 500 ms to 100 ms. Even still, 2 of the 11 (18.2%) subjects reported uncomfortable scalp sensations, and we aborted those experiments. Analysis of the other nine subjects’ data continued to find no statistical ipsilateral ROI power increases compared to the contralateral hemisphere, though subject average visualizations of the 30 ms ERP had a stronger concordance with the ROI than with the (on-average) lower target pressures used in our original study (Figure 3). The increased concordance provides some support for the argument that higher pressures may be more efficacious as a sole stimulus (as opposed to a co-modulator), though the balance of efficacy and the risk of scalp burns must always be weighed (especially given a report that the pressure levels needed for tFUS to generate direct suprathreshold stimulation in mice sciatic nerves lead to irreversible tissue damage^41^).

Why tFUS may be capable of stimulating the rodent cortex^9,40^, yet seems to be limited to subthreshold co-modulating the human cortex cannot be confirmed, but if we are to speculate, based on all that has been discussed thus far, the reason could be that the amount of ultrasound pressure loss through the skull in larger brain models does not reach the levels of cortical pressure needed to open enough piezo-electric channels. As the pressure is increased, so is the amplitude of the PRF beeping noise. Even if there is a small tFUS-induced targeted-location specific effect, it might be masked by the auditory evoked response. Another possibility could be that humans may not have as high a density of mechanosensitive channels for the ultrasound to activate as rodents.

There have been some studies, in humans^28,42^ and in non-human primates^43^, that indicate tFUS may have suprathreshold stimulation electrophysiological effects on deep-brain targets. This does, initially, seem contradictory to our results and theory. However, we postulate two reasons as to why these are actually complementary pieces of evidence. First, in rodents, the deep brain has been found to have a much higher concentration of piezo-electric ion channels than the cortex and is therefore more sensitive to ultrasound modulation^11^. In other words, for our HH model, G_piezo, deep brain_ >> G_piezo, cortex_, where G is the membrane conductance. In addition to having higher piezo-electric channel conductance, deep brain neurons have also been reported to have much lower activation thresholds than cortical ones^44^. As a result, what is subthreshold pressure on the cortex may be suprathreshold in the deep brain^28^. Second, despite targeting the motor cortex, we observed large scale activation in the auditory and deep brain structures for tFUS-only conditions at the 30, 90, and 170 ms peak windows (Figure 4a-b). This is not surprising, as the auditory pathways directly project into the deep brain. In fact, when we investigated connectivity from primary auditory cortex to the deep brain in our subjects, we found significantly increased A1 to deep brain Granger causality for both etFUS and itFUS (Figure 4c). Other research has demonstrated that frequencies beyond which humans can perceive as sound (20 Hz – 20 kHz) can still affect auditory pathways^45,46^, meaning that no ultrasound PRF is inherently exempt from this phenomenon. We corroborated this possibility with an additional follow up study with an inaudible (30 kHz) PRF and continued to find significantly increased A1 to deep brain Granger causality (Supplemental Figure S3). As a result, we deem it highly possible that deep brain regions can be “activated” by the auditory aspect of ultrasound, but then co-modulated from the pressure wave itself. In any case, more research into the piezo-electric properties of the human brain is warranted.

It is important to discuss the actual values of the localization scores beyond their statistical significance. In recent years, tFUS has garnered rising attention as a neuromodulation technology with millimeter precision^8,47,48^. Our *ex vivo* scans of the ultrasound’s pressure wave profile (Figure 1c) support this, but the average of our subject precisions of the tEAS reconstructed sources are only about 0.10 calculated by Equation 1 (Supplemental Figure S1b). This is not altogether unsurprising, as we applied modulation to resting-state brains. The resting-state brain is not ‘off’ in a traditional electronics sense, but rather unfocused. Some regions of the brain, known as the default mode network, are actually more active when a person is at rest^49^. The low precision scores may be partially attributed to heightened “off-target” activities.

The low scores may also be somewhat attributed to our choice of using Otsu’s method to determine the source threshold. Otsu’s method was first proposed for computer vision purposes to find a threshold value that minimized the intraclass variance (alternatively, it maximizes the interclass variance) between an image’s foreground and background^50^, and it has become a commonly used technique in EEG source localization to analytically and objectively determine the threshold between a source signal of the brain and noise in the recording^51^. For individual subject-level data, the thresholds were around 30% (etEAS range: 25 – 42%, itEAS range: 21 – 39%). Since brain tissue is conductive, any electrical activations induced by itEAS will propagate outwards until they spatially decay. These peripheral activations are still likely to be detected as sources given the low threshold values.

Even with persisting off-target activations, the increase in activity levels for the targeted area is apparent. The source-morphed subject averages (Figure 3a) as well as the grand average ERP (Supplemental Figure S9) converged to cover the ROI indicating the off-target activations are inconsistent across subjects compared to the stimulated area. Indeed, when we calculated the Euclidean distance from the grand average brain voxel with highest activity to the closest point in the ROI, both tEAS conditions had 0 mm localization error (compared to 73, 23, 73, and 26 mm for etFUS, itFUS, etDCS, and itDCS, respectively).

This is the first human study, to our knowledge, that harnesses and demonstrates the electro-acoustic compounding effect in neural stimulation. In doing so, we develop a new non-invasive technique that does not just modulate, but rather stimulates, targeted brain regions. Our results provide evidence that tFUS is a subthreshold co-modulator, and we have discussed how this theory is a unifying framework for, up until now, conflicting reports of tFUS neuromodulatory capabilities.

## 3. METHODS

27 healthy human subjects volunteered to participate in the study. Subjects took part in a pre-experiment session where they were safety-screened for magnetic resonance imaging (MRI) and tFUS eligibility, and they provided informed consent to continue with the study. The study protocol was approved by Advarra institutional review board. Subject recruitment and experiments were conducted in cohorts of 22 (subjects 101-122; 14 F / 8 M; mean ± std age: 24.5 ± 4.0 years old), 11 (subjects 201-211; 7 F / 4 M; mean ages = 23.22 ± 1.10 years old), and 11 (subjects 301-311; 5 F / 6 M; mean age = 24.2 ± 2.3 years old) participants. Some subjects participated in multiple cohorts of experimentation.

### MRI and MRI Segmentation

Subjects underwent a T1-weighted 3T Magnetic Resonance Imaging (MRI) structural scan of their brain at the CMU-Pitt BRIDGE Center (RRID:SCR_023356). The brain images were then segmented in FreeSurfer^52,53^ using the Desikan-Killiany atlas for post-hoc analysis. One subject was not MRI eligible due to an implanted device, so the FreeSurfer *FSAverage* brain was used^54^.

For EEG source imaging cortical parcellation, the primary auditory cortices and deep brain regions were defined by the left and right transversetemporal and medialwall parcellations, respectively. The IL/CL M1 ROI was defined by projecting the four IL/CL ROI electrodes orthogonally to the brain model’s inflated cortex and creating a square slice whose corners were defined by the electrodes.

### In silico transcranial focused ultrasound simulations

In the initial subject cohort, tFUS pressure was normalized with respect to the energy delivered on the scalp (0.98 MPa). To calculate the resulting on-cortex target pressure across subjects, Sim4Life (Version 7.0, Zurich Med Tech; Zurich, Switzerland)^55^ computer simulations of the single element tFUS transducer were conducted. Structural MRI files were converted to pseudo computed tomography (pCT) for use with the software^56,57^. For the simulations, the transducer was positioned over the subject’s left motor cortex, and the resultant pressure loss from brain tissue and skull bone attenuation was used to calculate spatial-peak temporal-average intensity (I_SPTA.3_; subject range = 48.6 – 372 mW/cm^2^), derated spatial-peak pulse-average intensity (I_SPPA.3_; 0.16 – 1.24 W/cm^2^), and Mechanical Index (MI_.3_; 0.10 – 0.28). The values for each subject are listed in Supplemental Table T1. In the case of two subjects, a pCT could not be generated (one subject did not have an MRI, and the other’s MRI had quality issues). All metrics were within the Food and Drug Administration’s guidelines for ultrasound (MI_.3_ < 1.9, I_SPPA.3_ < 190 W/cm^2^, and I_SPTA.3_ < 720 mW/cm^2^)^58^.

For later subject cohorts, Sim4Life simulations were used to normalize *in situ* tFUS pressure to 250 kPa and 300 kPa. The equivalent free water pressures are provided in the relevant supplemental data.

### Neuromodulation-EEG Experiment

Once the MRI segmentation was completed, subjects were asked to return for the neuromodulation-EEG portion of the study, which took place in an acoustic and electromagnetically shielded booth (IAC Acoustics; Naperville, Illinois, USA). Subjects were fit with a 64-channel EEG cap (BrainCap TMS, Brain Products, Gilching, Germany). EEG caps were aligned with subject landmarks following standard convention (centering electrode Cz between both the inion and nasion, as well as the right pre-auricular and left pre-auricular). EEG electrode impedances were lowered to be less than 10 kΩ using ABRALYT HiCl electrolyte gel.

The 4x1 high-definition tDCS (Soterix Medical Inc., Woodbridge, NJ, USA) electrodes were placed in the following locations: 1 peripheral electrode between aFz (ground), Fz, F1, and AF3, 1 peripheral electrode between CP1, CPz, Pz, and P1, 1 peripheral electrode between FC3, C3, C5, and FC5, 1 peripheral electrode between FC2, FC4, C2, and C4, and the center electrode between FCz (reference), Cz, C1, and FC1 (Figure 1b). The electrical dosage for each subject was based on their own personal threshold for feeling scalp sensations. To determine the dosage, current was slowly ramped up from 0 until the subjects reported sensations. The dosage was set at 0.1 mA below their perception threshold, up to a maximum of 2 mA (Supplemental Table T1). The same amperage was used across all conditions involving tDCS (etEAS, itEAS, etDCS, and itDCS).

Excitatory er-anode current was defined as inhibitory tDCS current. tDCS remained on throughout for the entirety of the 200 trials per condition.

For on-target ultrasound conditions (etEAS, itEAS, etFUS, and itFUS), a 500 kHz f_0_ (fundamental frequency) single-element ultrasound transducer (Blatek Industries Inc., Boalsburg, PA, USA) was attached between EEG electrodes C1, C3, FC1, and FC3. Using an image-guided neuronavigation system (MagVenture Localite TMS Navigator; Alpharetta, Georgia, USA), the ultrasound target was visually confirmed to align with the subject’s left motor cortex. For off-target ultrasound spatial control conditions (etDCS, itDCS, and an additional 3 kHz tFUS), the ultrasound was targeted to the subject’s right prefrontal cortex area between electrodes AF4, AF8, F4, and F6. Based on our lab’s prior work, excitatory ultrasound was defined by a PRF of 3 kHz, and inhibitory ultrasound by a PRF of 30 Hz^59^ (Figure 1b). The duty cycle (30%) and sonication duration (500 ms) were held constant.

For each condition, subjects were asked to assume a resting state for 200 trials (Figure 1d). Specifically, they were instructed to close their eyes, remain as still as possible, and to relax their thoughts in a meditative-like state. The inter-trial interval was 2 seconds with up to a 20% (0.4 s) random jitter. A five-minute break was taken between each condition to allow for the subjects to stretch and let any cumulative effects of neuromodulation dissipate. Experimental condition order was randomized across subjects, though data for e/itDCS control conditions were not collected in the initial subjects (Figure 2d).

One follow up study was conducted to investigate additional duty cycles (0.6%, 10%, and 60%) for 30 Hz and 3 kHz PRFs, and additional pulse repetition frequencies (1500 Hz and 40 Hz) with 30% DC. Changing tEAS parameters was also done in this follow up to determine if the tEAS response differed as a function of PRF, but these conditions were always tested after the resting-state tFUS ones in order to minimize possible long-term contamination effects of tDCS. Another follow up session was done to investigate the effects of normalizing in-situ brain pressure. Other than these parameter changes, the protocol remained the same.

### EEG Analysis

#### EEG Preprocessing

EEG data were bandpass filtered between 1 and 40 Hz with a zero-phase infinite impulse response filter. Data were epoched from 500 ms before the trial onset time to 1.5 seconds after. Bad channels were detected by averaging all the trials and identifying double median absolute deviation (double MAD; threshold = 3) outlier pre-stimulus variance or total trial peak-to-peak amplitude. On the un-averaged trials, independent component analysis eyeblink and muscle artifacts were removed using MNE Python’s^60^ automatic artifact detection with a z-threshold of 2.5. The data were common average referenced, and the previously identified bad channels were interpolated back into the signal using MNE Python’s interpolate_bads function. The cleaned data were averaged over all the trials to get the evoked waveform and z-scored with respect to the -500 to -100 ms pre-stimulus window. In a few instances, active tDCS created noticeable EEG artifacts. These data were excluded in the analysis (Figure 2d).

#### EEG Sensor Domain Analysis

Preprocessed EEG waveforms were averaged across subjects for each condition and used to produce butterfly plots (Figure 2a). Statistical analysis was conducted using non-parametric spatio-temporal two-tailed cluster tests (5,000 permutations, α = 0.05; Figure 2b). Paired t-tests were conducted comparing ipsilateral ROI electrodes C1, C3, FC1, and FC3 against their homologous contralateral counterparts C2, C4, FC2, and FC4 within the exogenous (0 – 200 ms^34^) brain response window. Significantly different adjacent data were clustered together in time and space. Data were randomly shuffled and the largest significant cluster from each permutation was used to construct a cluster-size histogram. The cluster-forming threshold was set at a significance level of 0.02. Clusters from the original data with sizes greater than the α percentile (corresponding to *p* < 0.05) were marked as significant.

ROI polarity analysis for largest overlapping significant cluster components of tEAS was determined by taking the sign of electrode C3’s average over the response window (10 – 63 ms). McNemar’s test, a statistical procedure for assessing paired binary data, was used to investigate the effect of excitatory and inhibitory modulation parameters on ROI polarity for each condition. False discovery rate (FDR) multiple comparison correction was applied.

#### EEG Source Imaging

EEG source imaging (ESI) was solved by minimizing the difference between cortical current density produced scalp EEG and recorded EEG^4^ using MNE Python’s implementation of dSPM, which normalizes the minimum norm estimation relative to a noise covariance matrix^61^. The noise covariance used for dSPM was calculated from each condition’s baseline window (-500 to -100 ms). EEG electrodes were anatomically aligned to the subject’s specific MRI model using the standard montage setup. The dSPM source maps were averaged over the 10 – 63 ms response window for ROI activity and localization analysis. Subject source maps were morphed from their individual head models to FreeSurfer’s *FSAverage*^54^ brain model for group-level visualizations (taking the average of each subject’s max-min scaled data) and connectivity analysis. For grand average ERP analysis (Supplemental Figure S9), the subjects’ EEG data were averaged prior to EEG source imaging with the *FSAverage* brain model.

For each subject, the mean activity, precision (Equation 1), recall (Equation 2), and F1 score (Equation 3) were calculated for the source-imaged activity with respect to the ipsilateral ROI. The ROI was defined by all the cortical space within a bounding box formed by projecting electrodes C1, C3, FC1, and FC3 (the electrodes between which tFUS was targeted) orthogonally from the scalp to the cortex. For precision, recall, and F1 score, *true* or *false positives* were defined as active brain sources that were inside, or outside, respectively, the region of interest*. False negatives* were defined as brain voxels within the region of interest that were not estimated as active sources. The threshold for what constituted a brain source, compared to noise, was defined using Otsu’s method^50,51^. Pairwise statistical comparisons between the mean activity, precision, recall, and F1 score were conducted with linear mixed effect modeling (see *Linear Mixed Effect Modeling* for more details.).

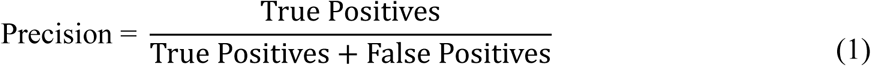

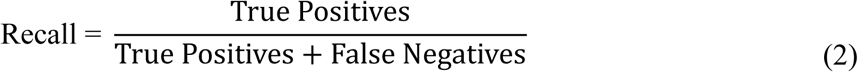

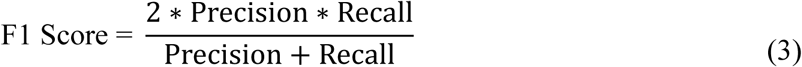

The contralateral ROI was defined by the cortical space contained within the projection of electrodes C2, C4, FC2, and FC4. Comparisons between ipsilateral and contralateral mean activity were conducted with 5,000 permutation one-tailed paired t-tests and FDR multiple comparison correction (H_1_: μ_IL_ > μ_CL_). A permutation t-test is a nonparametric alternative to the parametric t-test, which relies on no underlying assumptions about data distributions. Instead, it shuffles the data between conditions an *N_permutations_* number of times (in this case 5,000) and calculates the T-statistic for each instance. This generates a histogram of T-statistics, which the unshuffled data’s T-statistic is compared against to determine its significance. The *p*-value represents the proportion of permuted T-statistics that are at least as extreme as the unpermuted T-statistic. In some instances, comparisons across the ipsilateral activity for multiple conditions were assessed with permutation ANOVA tests. The permutation ANOVA test follows the same logic as the permutation t-test, except is it based on the F-statistic. Bayes Factor ANOVA testing was performed in R (version 4.3.3) with the BayesFactor package.

#### Connectivity Analysis

Changes in Granger causality were calculated on source waveforms^62^ for the primary auditory cortex to deep brain and the targeted ROI to deep brain. Granger causality was calculated using the Toda and Yamamoto procedure^63^. The lag for each pathway was selected using the Akaike Information Criterion with possible choices between 10 and 15 ms based on reported values for transmission delays^64^. Net Granger causality (GC_Net_) was calculated by subtracting the reverse Y ➔ X Granger causality Wald statistic from the forward X ➔ Y Granger causality Wald statistic. GC_Net_ was calculated for each 200 ms time bin from stimulus onset to one second after, as well as for the -0.5 to -0.1 s pre-stimulus baseline period. Permutation paired two-tailed t-tests were used to compare each time bin’s GC_Net_ to the pre-stimulus period. FDR multiple comparison correction was applied locally for each series.

Spectral connectivity was assessed with imaginary coherence (Equation 4), a metric used for quantifying non-zero time lagged undirected connectivity^65^. The power spectral density (PSD) for each series was calculated using Welch’s method with a 0.2 s window in 0.2 s bins over the 0 to 1 second post-stimulus period and mean-normalized against the -0.5 to -0.1 s baseline period. Imaginary coherence was calculated for alpha (10 – 13 Hz), beta (13 – 30 Hz), and gamma (30 – 40 Hz) frequency bands. Comparison between the ipsilateral and contralateral changes in ImCoh for each time bin were assessed with 5,000 permutation t-tests. FDR multiple comparison correction was applied locally for each condition/frequency band level.

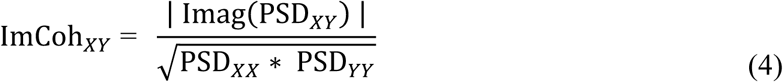

#### Linear Mixed Effect Modeling

The partially-paired data (Figure 2d) made linear mixed effect models excellent candidates for analyzing statistical differences across conditions. Linear mixed effect models account for both fixed effects (experimental conditions) and random effects (subject-to-subject differences) and estimate the true mean of the predictors on the outcome variables. For all our analysis, we modeled the relationship as Equation 5. The outcome variable for condition *j* and subject *k* is a function of some fixed offset for all subjects and all conditions, 𝛽_0_ , the estimated true mean effect of experimental condition *j*, 𝛽_1,j_, and a subject-specific random effect for subject *k*, *b_k_*. The difference between the model’s prediction and the observed outcome is the error term 𝜖_j,𝑘_ . Since our independent variables are categorical, 𝛽_1_ is a (1 x J) vector of the estimated true means of the effects, and *Condition* is a (J x 1) one-hot vector. The *outcome* variable is a stand-in for mean ROI activity, precision, recall, or F1 score. Modeling was done in R (version 4.3.3) with the lme4 and multcomp packages.

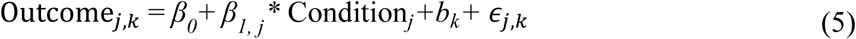

Pairwise comparisons were made using R’s glht function, which calculates the mean difference of the estimated true effect across conditions and their pooled standard error. These values are used in *z*-tests to statistically compare condition means with FDR multiple comparison correction. The pooled standard error, used in calculating Cohen’s *d* (Equation 6), was found by multiplying the pooled standard error by the square root of the subject number.

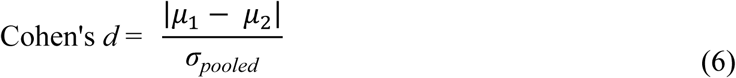

The primary assumptions of a linear mixed effect model are: 1) the response variable is linearly related to the predictor variable(s), 2) the model errors are independent, 3) the error variances are homogeneous, and 4) the errors are normally distributed. Since the independent variables are categorical (i.e., experimental condition), the model is calculating group means and not fitting a line, so assumption 1 is automatically satisfied. Non-violation of assumption 2 was tested by plotting the autocorrelation of the model residuals and identifying no significant systematic patterns. Assumptions 3 and 4 were assessed with Levene and Shapiro-Wilks tests, respectively. To meet the requirements of assumptions 3 and 4, ROI activity (strictly positive) were natural log transformed, and classification metric data (precision, recall, and F1; inclusively bound between 0 and 1) were square-root transformed.

### Hodgkin Huxley Modeling

To model the electro-acoustic effect, the Hodgkin Huxley model was modified to include a piezo-electric channel. Standard model parameters from the original Hodgkin Huxley paper were used for the sodium, potassium, and leak channels^35^. Piezo-electric channels were modeled as an additional parallel pathway within the circuit (Figure 6a; Equation 7). In mice, the reversal potential of the piezo-electric ions (*E_piezo_*) has been reported to be 0 mV^36^, single-channel conductance are approximately 26.6 pS^37^, and piezo-electric channel densities have been calculated as about 1 channels/µm^2^ ^38^. Converting these to model terms, which reference the resting membrane potential at 0 mV and not -65mV, we approximated *E_piezo_* as +65 mV and *G_piezo_* as 2.65 mS/cm^2^. The last component for modeling this pathway of the circuit is to consider the probability that the piezo-electric channels are open. Outside of pressure ranges where the channel has a 0 or 1 probability of being open, the open probability (*P_open_*) scales approximately linearly with applied pressure^38^. As a result, *P_open_* was modeled directly as the ultrasound waveform (clipped at 0 pressure to avoid negative probabilities), scaled to have some peak probability less than 1.

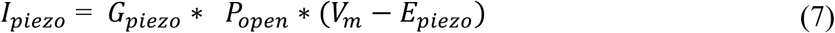

### Data availability statement

De-identified human data will be shared via Figshare upon paper acceptance.

### Code availability statement

Customized code will be shared via GitHub upon paper acceptance.

### Authorship contribution statement

Conceptualization: J.K., K.Y., and B.H. Methodology: J.K., C.G., K.Y., and B.H. Investigation: J.K., C.G., K.Y., and B.H. Data curation: J.K., J.Z., Y.D., and Y.Z. Formal analysis: J.K., C.G. Visualization: J.K. Writing—original draft: J.K. Writing—reviewing and editing: J.K., C.G., K.Y., and B.H. Supervision: B.H.

## Acknowledgements

This work was supported in part by NIH grants R01NS124564 (PI: B.H.), RF1NS131069 (PI: B.H.), R01NS127849 (PI: B.H.), R01NS096761 (PI: B.H.), and T32EB029365 (J.K. and C.G.; PI: B.H.). J.K.’s work was supported in part by the National Science Foundation Graduate Research Fellowship Program under Grant No. DGE2140739. Any opinions, findings, and conclusions or recommendations expressed in this material are those of the authors and do not necessarily reflect the views of the National Institutes of Health and the National Science Foundation.

The authors would also like to thank Kings Jiang for her assistance in EEG capping, Elena Bondi, Jesse Rong, Chih-Yu Yeh, and Annabel Frake for discussions on EEG analysis and preprocessing, Yunruo Ni and Chih-Yu Yeh for assistance in 3D printing tFUS transducer holders, and Zherui Li for assistance in converting subject MRIs into pseudo-CT images. In silico tFUS simulations were conducted using Sim4Life by ZMT, www.zmt.swiss.

## Competing Interests Statement

B.H., K.Y., and J.K. are co-inventors of pending patent applications.

