## Supplemental Information for "Transcranial focused ultrasound induces source localizable cortical activation in resting state humans when applied concurrently with transcranial electric stimulation"

### SUPPLEMENTAL MATERIALS

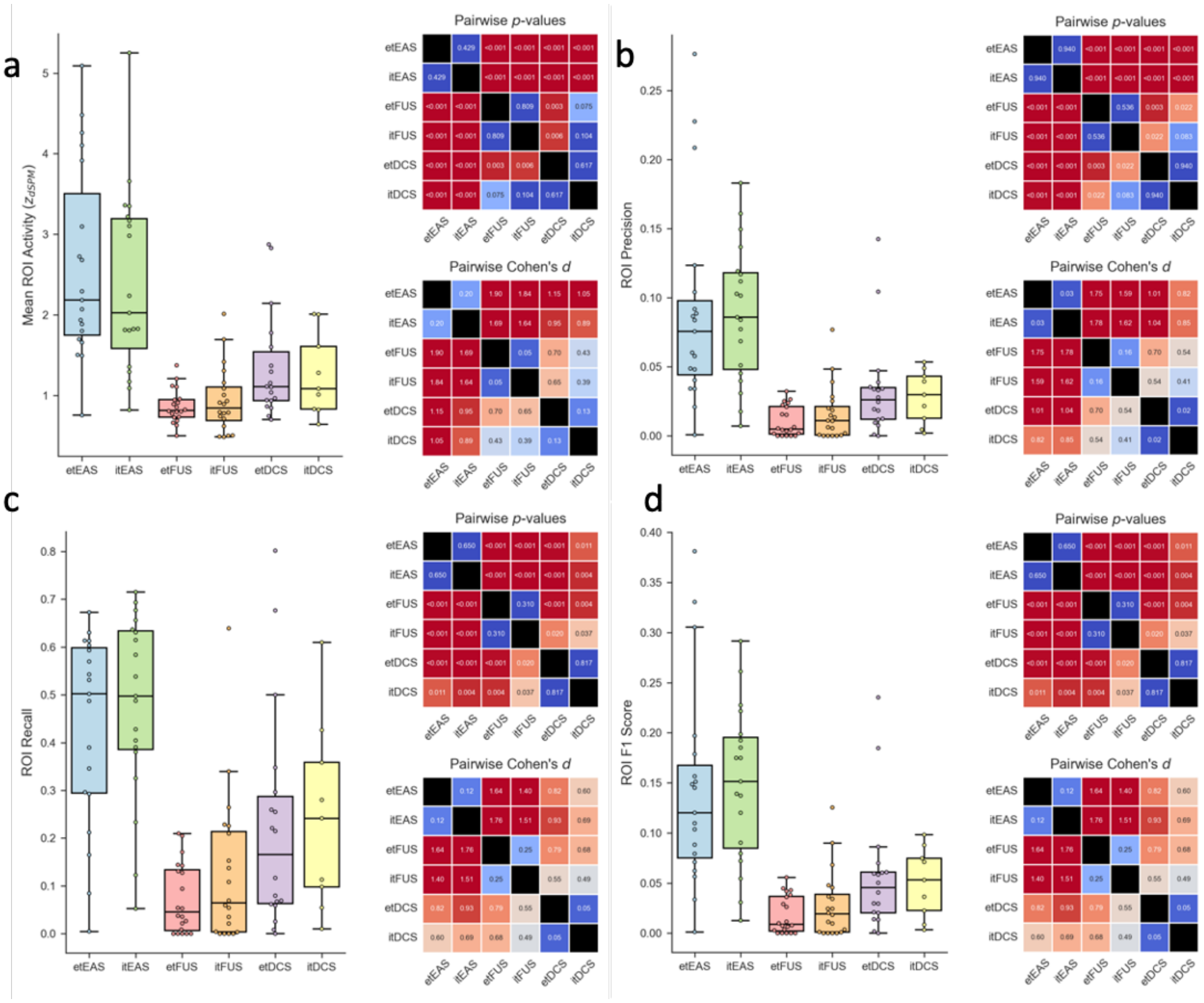


**Supplemental Figure S1: tEAS induces significantly more localizable cortical responses than tFUS or tDCS.** Pairwise testing for a) mean ipsilateral source power, b) ipsilateral precision, c) ipsilateral recall, and d) ipsilateral F1 score were conducting using linear mixed effect models. For localization metrics (precision, recall, and F1 score), brain source “positives” were those with activity higher than Ostu’s threshold for given dataset. For linear mixed-effect modeling, ROI power (strictly positive) were log transformed and localization metrics (bound between 0 and 1) were square-root transformed. Model results yielded e/itEAS induced significantly higher ipsilateral ROI source power, ROI precision, ROI recall, and ROI F1 score than the other conditions (two-tailed z-tests with FDR multiple comparison correction). Cohen’s *d* analysis indicates that for all pairwise comparisons with tFUS or tDCS, tEAS induced at least a moderately sized effect ($d > 0.5$).

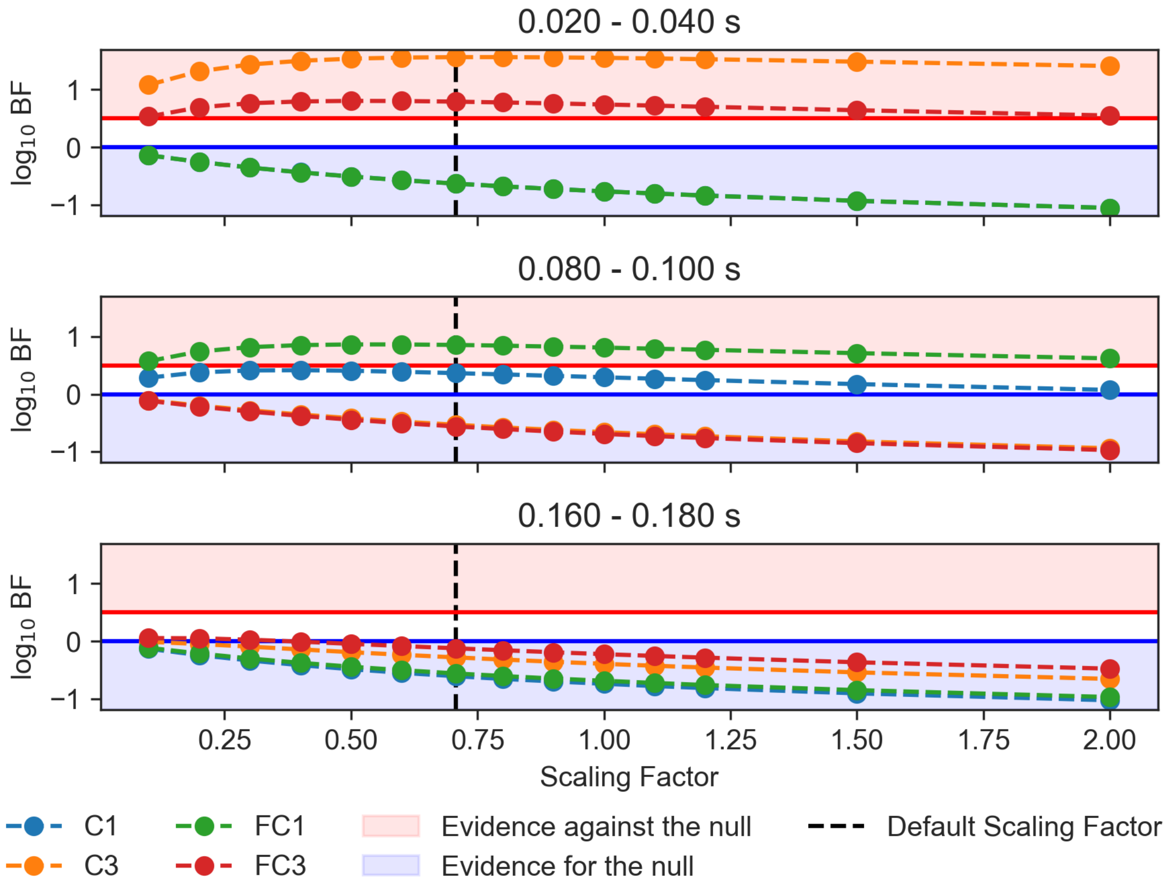


**Supplemental Figure S2**: **Bayes Factor Analysis indicates the presence of robust significant differences between etEAS and etFUS conditions in the first two AEP ERP time windows**. Bayes Factor t-tests were run on average activity for the specified electrodes in each time window and compared between conditions. In the 20 – 40 ms time window, there is robust evidence that the response of electrodes C3 and FC3 were significantly different across conditions. In the 80 – 100 ms window, there is evidence that the response of FC1 differed.

**a**

| Subject | Pressure Ratio | In Situ Pressure | Free Water Pressure |
|---|---|---|---|
| 201 | 26.50% | 250 kPa | 943 kPa |
| 202 | 25.20% | 250 kPa | 992 kPa |
| 203 | 20.02% | 200 kPa | 999 kPa |
| 204 | 31.00% | 250 kPa | 806 kPa |
| 205 | 28.46% | 250 kPa | 879 kPa |
| 206 | 23.40% | 200 kPa | 855 kPa |
| 207 | 36.30% | 250 kPa | 689 kPa |
| 208 | 23.23% | 200 kPa | 861 kPa |
| 209 | 38.50% | 250 kPa | 649 kPa |
| 210 | 26.26% | 200 kPa | 761 kPa |
| 211 | 18.79% | 180 kPa | 958 kPa |

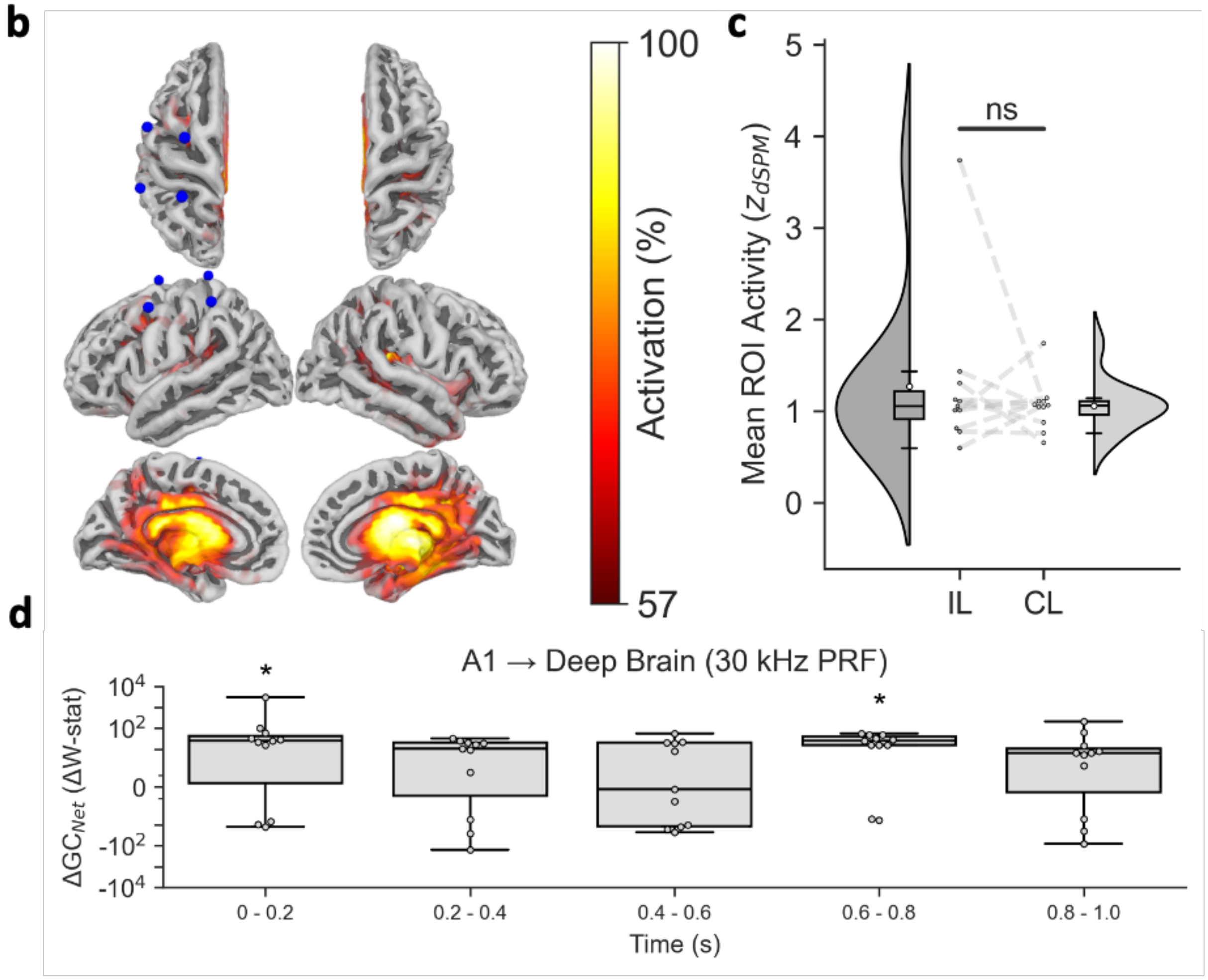


**Supplemental Figure S3**: **30 kHz PRF continues to induce net auditory to deep brain causality.** A follow up study was conducted with 11 subjects to investigate possible pathway auditory activations in an inaudible PRF range. The target pressure was normalized to 250 kPa using Sim4Life simulations, with a duty cycle of 30% and a sonication duration of 500 ms. Three subjects reported thermal discomfort during at 250 kPa in situ pressure, and their experiments were aborted and restarted with 200 kPa in situ pressure. Subject 211's pressure pre-emptively lowered to 180 kPa due to observations that > 1 kPa free water pressure were leading to heating in prior subjects with these tFUS parameters. **a)** Pressure ratios and levels are presented. **b)** The subject-average cortical response in the 10 – 63 ms window continues to show strong deep brain and temporal lobe activations. **c)** Comparisons of ipsilateral vs. contralateral ROI continue to yield no significantly increased ipsilateral power. **d)** Net Granger causality analysis was run for each subject to assess the degree to which primary auditory cortex (A1; FreeSurfer's transversetemporal parcellation) forecasted their deep brain activity (FreeSurfer's Medial Wall parcellation). FDR multiple comparison correction was applied. Inaudible tFUS (30 kHz PRF). Despite 30 kHz PRF being above the range of human hearing, significant increases in the net flow of information from auditory cortex to deep brain were observed. * $p_{adj} < 0.05$

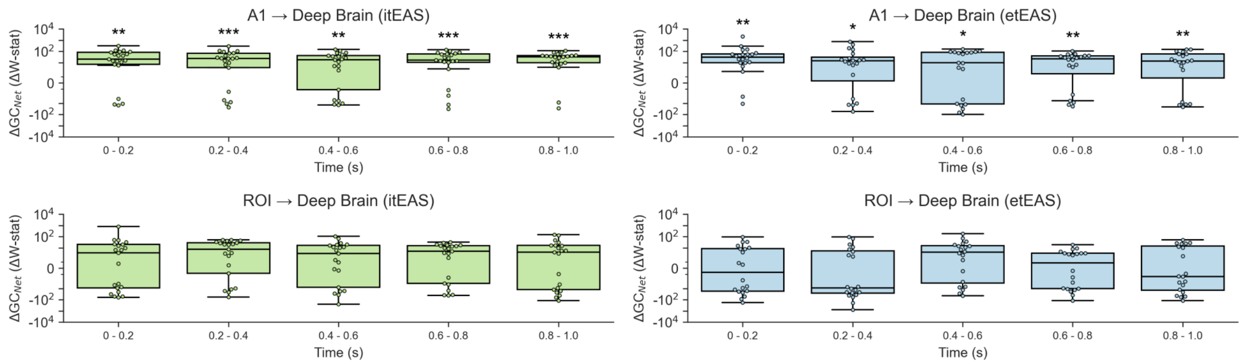


**Supplemental Figure S4**: **Auditory confounds persist in tEAS conditions.** Transcranial electroacoustic stimulation does not induce increase information flow from the targeted ROI (left M1) to the deep brain. The Net Granger Causality ($GC_{Net}$) Wald statistic was calculated for each time bin by subtracting the GC of the reverse flow from the GC of the forward flow of information. The $GC_{Net}$ values were normalized by subtracting the $GC_{Net}$ for the baseline period (-0.5 to -0.1 s) for each subject's pathways ($\Delta GC_{Net}$). The differences between the baseline $GC_{Net}$ and each time bin's $GC_{Net}$ were analyzed with permutation t-tests. FDR multiple comparison correction was applied locally. Both etEAS and itEAS continue to reflect strong auditory projections from A1 to the deep brain, and neither etEAS nor itEAS (despite producing M1 source localizable signals) induced significant M1 to deep brain Granger causality. *$p_{adj} < 0.05$, **$p_{adj} < 0.01$, ***$p_{adj} < 0.001$

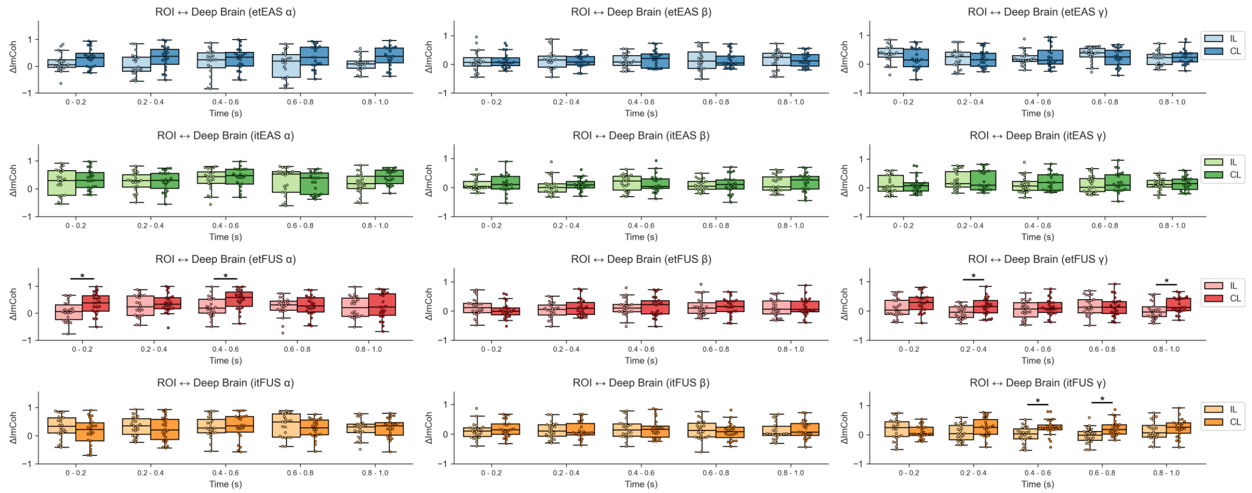


**Supplemental Figure S5**: **Despite not significantly altering the directed *net flow* of information from M1 to deep brain, tFUS alters the left M1 ROI to deep brain spectral connectivity.** As opposed to net Granger causality's measure of directed signal flow, imaginary coherence (ImCoh) can be used to quantify a more general undirected spectral connectivity. The absolute ImCoh was calculated for spectral frequencies α (10 – 13 Hz) β (13 – 30 Hz), and γ (30 – 40 Hz) for the 0 to 1 s post-stimulus period using Welch's method with 0.2 s windows . The minimum frequency (10 Hz) was chosen as to allow for two full cycles in the 0.2 s window. ImCoh were mean corrected (ΔImCoh) by subtracting the average of each condition's pre-stimulus -0.5 to -0.1 s window. Differences between ipsilateral and contralateral ΔImCoh were assessed with permutation t-tests. FDR multiple comparison correction was applied locally for each condition/frequency band of interest. Both tFUS conditions significantly reduced endogenous (> 200 ms) gamma band connectivity. etFUS also significantly reduced alpha hand spectral connectivity. None of these differences were present in the tEAS conditions, indicating that the suprathreshold electro-acoustic effect reconnected these dissociations. *$p_{adj} < 0.05$

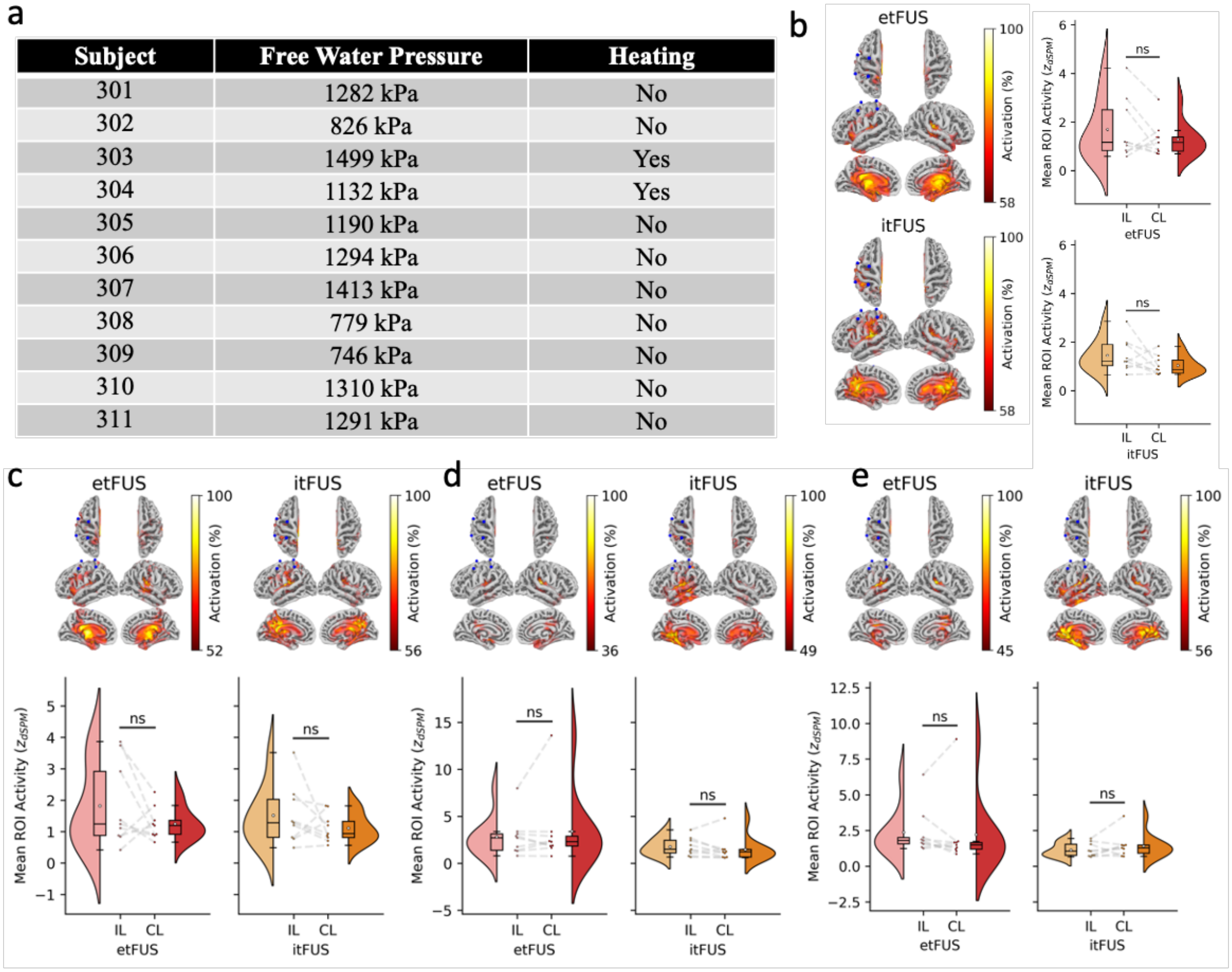

| Subject | Free Water Pressure | Heating |
|---|---|---|
| 301 | 1282 kPa | No |
| 302 | 826 kPa | No |
| 303 | 1499 kPa | Yes |
| 304 | 1132 kPa | Yes |
| 305 | 1190 kPa | No |
| 306 | 1294 kPa | No |
| 307 | 1413 kPa | No |
| 308 | 779 kPa | No |
| 309 | 746 kPa | No |
| 310 | 1310 kPa | No |
| 311 | 1291 kPa | No |



**Supplemental Figure S6: Higher pressure did not result in source localizable activity.** The 3 kHz and 30 Hz PRF resting state tFUS experiments were repeated on a cohort of eleven healthy human subjects, this time normalizing the in situ pressure to 300 kPa ($I_{spta.3}$ = 217 mW/cm$^2$, $I_{sppa.3}$ = 72.2 W/cm$^2$, $MI_{.3}$ = 0.212). To reduce heat build up at the interface of the scalp and and transducer, the sonication duration was reduced to 100 ms. Despite this precaution, two of the eleven subjects (18.2%) still reported uncomfortable scalp heating from the ultrasound transducer and their experiments were aborted. Their data are not included in group-level analysis. **a)** Free water equivalent pressure levels and whether or not heating occurred are presented. **b-d)** Subject average visualizations and ipsilateral vs. contralateral ROI activity are provided for **a)** 10 – 63 ms **b)** 20 – 40 ms, **c)** 80 - 100 ms, and **d)** 160 - 180 ms ERP windows. With the increased pressure, subject average visualizations did appear slightly more concordant with the ROI within the 10 – 63 and 20 - 40 ms time windows. However, statistical comparisons (one-tailed exact permutation paired t-test with FDR multiple comparison correction applied for each peak window) remained non-significant for all comparisons (all $p_{adj} > 0.05$).

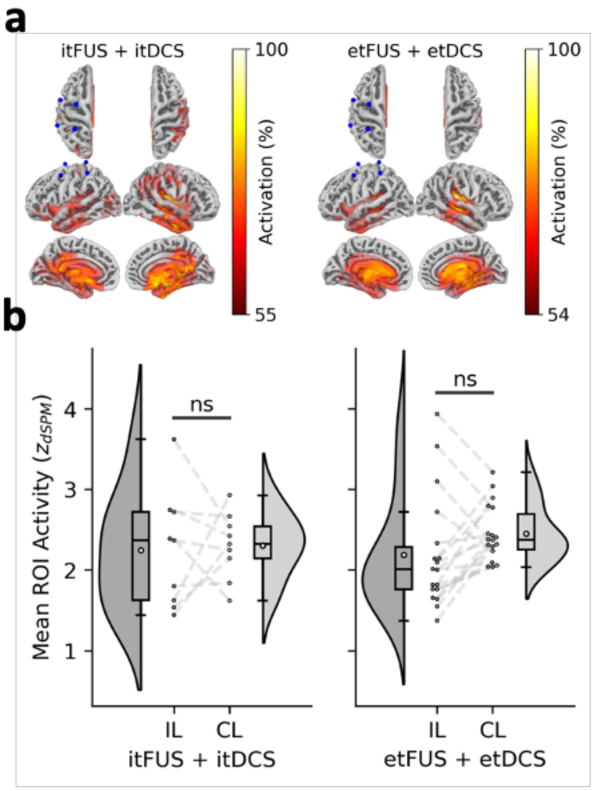


**Supplemental Figure S7**: **The transcranial electro-acoustic effect is more than just the summation of independent electric and acoustic modulation. a)** Source-imaging data for itFUS and itDCS, as well as etFUS and tDCS, were added together for each subject, and then averaged across subjects. Visualizations continue to show strong off-targeted sources. **b)** The average ipsilateral ROI activity of the added sources continues to be not significantly greater than the contralateral ROI for both the summed excitatory and summed inhibitory conditions ($p > 0.05$).

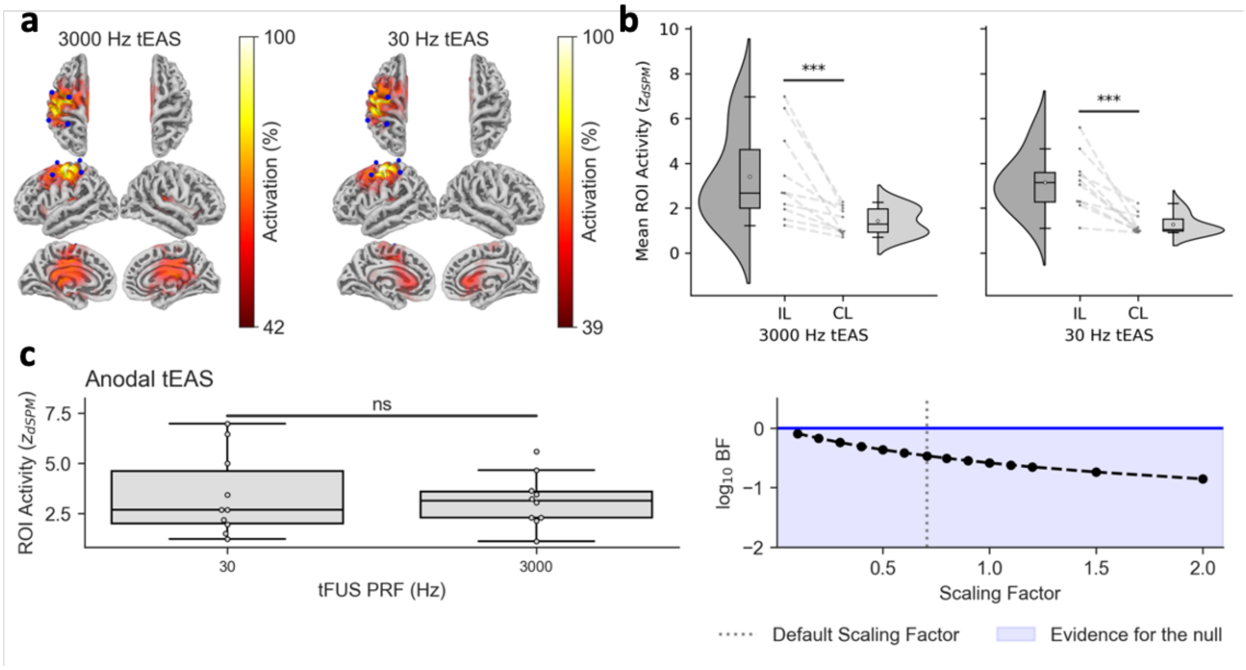


**Supplemental Figure S8**: **The electroacoustic response is not driven by PRF.** A follow up study was conducted on 10 subjects where the same tDCS polarity (anodal) was combined with 30 Hz and 3 kHz PRF tFUS. **a)** Data were back projected to their cortical space and the brain sources for tEAS continue to be concordant with the ROI. **b)** Comparison of Ipsilateral vs. Contralateral ROI source activities show both 30 Hz and 3 kHz PRF tEAS produce significantly stronger ipsilateral than contralateral responses. **c**) Comparison between the ipsilateral response for 30 Hz tEAS and 3 kHz tEAS reveal no significant difference between the two. ***$p_{adj}$ < 0.001

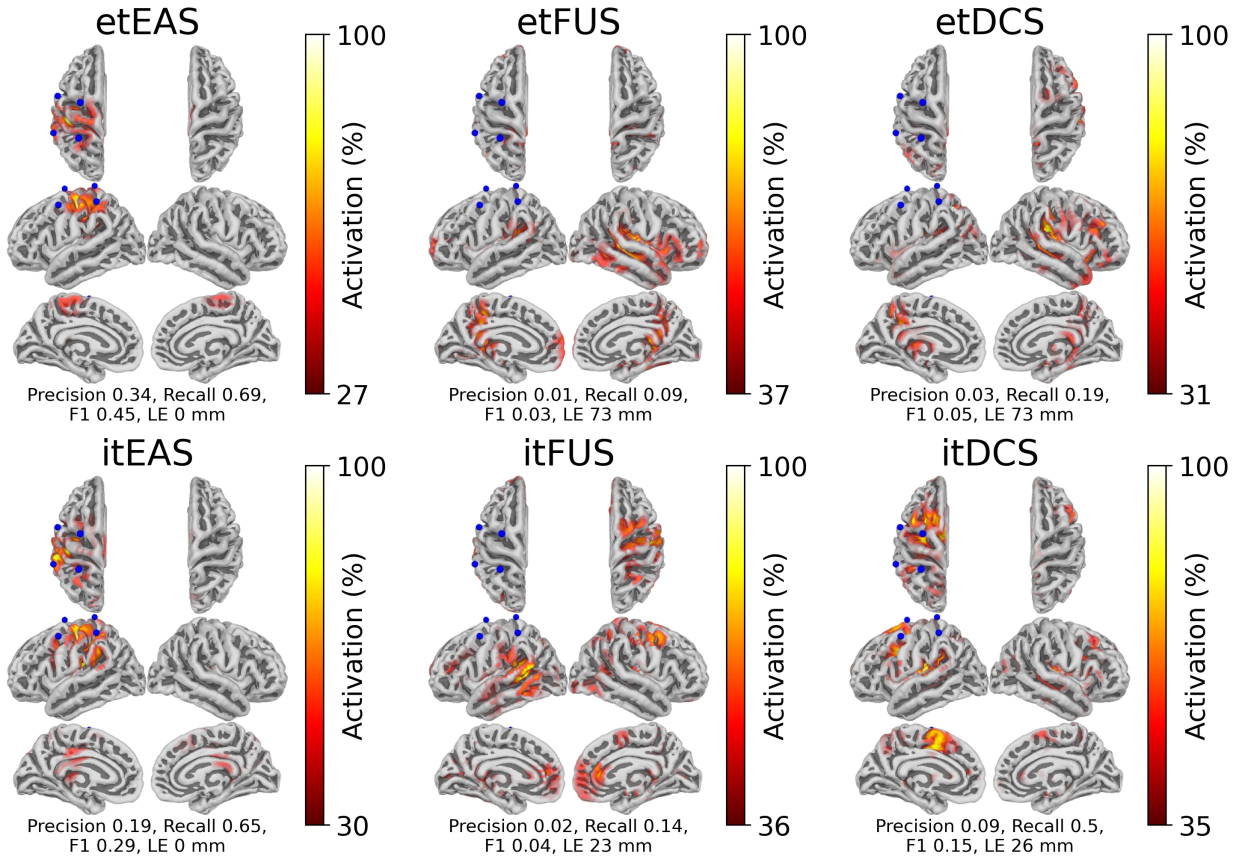


**Supplemental Figure S9**: **Grand Average source localization results are consistent with the averaged subject-specific source localizations.** Subject EEG data were averaged across trials and z-scored with respect to the -0.500 to -0.100 s window. Then, subject data were averaged together to form a grand average ERP. EEG source imaging was performed using the grand average ERP and the *FSAverage* head model. Results are consistent with those obtained by performing source localization of subject data first, source morphing to the common template, and then averaging (Figure 3). Brain source activity thresholds were determined using Otsu's method. Localization error was computed as the Euclidean distance from the brain voxel with highest activity to the closest point in the ROI.

### Supplemental Table T1: Subject-specific Neuromodulation Dosage

| Subject ID | $Isppa_{.3}$ (W/cm$^2$) | $Ispta_{.3}$ (mW/cm$^2$) | $MI_{.3}$ | Current (mA) |
|---|---|---|---|---|
| **Subj101** | 0.22 | 66.89 | 0.12 | 2.0 |
| **Subj102** | 0.21 | 61.56 | 0.11 | 1.5 |
| **Subj103** | N/A (Poor quality MRI could not be converted to pCT) | | | 1.5 |
| **Subj104** | 1.02 | 306.08 | 0.25 | 2.0 |
| **Subj105** | 0.16 | 48.57 | 0.10 | 1.5 |
| **Subj106** | 0.18 | 54.80 | 0.11 | 1.3 |
| **Subj107** | 0.21 | 62.32 | 0.11 | 0.7 |
| **Subj108** | 0.22 | 65.95 | 0.12 | 1.5 |
| **Subj109** | 0.42 | 125.35 | 0.16 | 1.1 |
| **Subj110** | 0.31 | 93.10 | 0.14 | 1.0 |
| **Subj111** | 0.74 | 223.22 | 0.22 | 1.0 |
| **Subj112** | 0.27 | 82.01 | 0.13 | 1.3 |
| **Subj113** | 0.23 | 68.96 | 0.12 | 0.9 |
| **Subj114** | 0.54 | 163.12 | 0.18 | 2.0 |
| **Subj115** | 1.24 | 372.02 | 0.28 | 1.4 |
| **Subj116** | 1.15 | 344.30 | 0.27 | 2.0 |
| **Subj117** | N/A (Ineligible for MRI) | | | 1.0 |
| **Subj118** | 1.00 | 299.37 | 0.25 | 1.4 |
| **Subj119** | 0.33 | 99.82 | 0.14 | 1.1 |
| **Subj120** | 0.34 | 101.17 | 0.14 | 0.7 |
| **Subj121** | 0.34 | 102.05 | 0.15 | 1.4 |
| **Subj122** | 0.47 | 139.77 | 0.17 | 1.0 |
